# Integrated ambient modeling and genetic demultiplexing of single-cell RNA+ATAC multiome experiments with Ambimux

**DOI:** 10.1101/2025.08.21.671671

**Authors:** Marcus Alvarez, Terence Li, Seung Hyuk T. Lee, Uma Thanigai Arasu, Ilakya Selvarajan, Tiit Örd, Elior Rahmani, Zeyuan (Johnson) Chen, Oren Avram, Asha Kar, Dorota Kaminska, Ville Männistö, Eran Halperin, Jussi Pihlajamäki, Chongyuan Luo, Minna U. Kaikkonen, Noah Zaitlen, Päivi Pajukanta

## Abstract

Single cell technologies have advanced at a rapid pace, providing assays for various molecular phenotypes. Droplet-based single cell technologies, particularly those based on nuclei isolation, such as simultaneous RNA+ATAC single-cell multiome, are susceptible to exogenous ambient molecule contamination, which can increase noise in cell type-level associations. We reasoned that genotype-based sample multiplexing can provide an opportunity to infer this ambient contamination by leveraging DNA variation in sequenced reads. Thus, we developed ambimux, a likelihood-based method to estimate ambient fractions and demultiplex single-cell multiome experiments using genotype-level data. Ambimux models the ambient or nuclear probability at the read level and thus can classify empty droplets and estimate droplet-specific ambient molecule fractions in each modality. We first evaluated our method using simulated data sets across a range of parameters. We found that ambimux closely estimated the ground truth droplet contamination fractions in the RNA (MAE=0.048) and ATAC (MAE=0.042) modalities. As a result, ambimux maintained high specificity (>95%) and was able to correctly assign singlets at considerably high ambient fractions (up to 60%) for both RNA and ATAC modalities. In comparison with models that do not consider ambient contamination, these only maintained similar sensitivity levels at considerably lower ambient fractions (up to 25%). We then generated a real data set of seven visceral adipose tissue biopsies run on a single 10x Multiome channel. We ran ambimux and detected 4,986 singlets, capturing similar numbers as other methods.

Then, we sought to evaluate the fidelity of the ambient fraction estimates from ambimux. We split singlets into ambient-enriched (>5% contamination in both modalities) or nuclear-enriched (<5% in both) droplets and performed gene-peak linkage analysis. Low ambient droplets resulted in more significant hits with gene-peak links enriched at the transcription start site relative to high ambient droplets, suggesting that the ambient droplets identified by ambimux hamper the identification of biologically meaningful signals. In summary, we developed a joint single-cell multiome demultiplexing method, ambimux, that accurately models and estimates ambient molecule contamination in each modality.

## Introduction

The ability to profile multiple molecular phenotypes in single cells has provided powerful opportunities to study the interaction between regulatory elements and gene expression^1,2^. Single cell technologies can profile molecular phenotypes beyond gene expression, including open chromatin^3^, methylation^4^, histone modifications^5^, and chromatin configuration^6^. These assays allow for the characterization of *cis*-regulatory elements (CREs) that regulate gene expression in a cell-type-specific manner^7^. These analyses are further aided by computational methods that can integrate separate single-cell ATAC-seq and RNA-seq experiments^8^. However, these approaches only infer the joint epigenomic and transcriptomic profiles of cell types without direct measurements in the same cell. To overcome these challenges, so-called multiome technologies have been developed to jointly profile open chromatin and RNA in the same cell. This can provide valuable insights in connecting CREs with gene expression, particularly in differentiating cell types where the epigenetic state may temporally mismatch gene expression^9^.

While joint profiling of single cells provides clear advantages over single-modality assays, it is currently costly and thus difficult to scale over many samples^10^. These limitations may hinder population-scale studies of gene regulation at the single-cell level. Multiplexing by pooling samples provides a feasible approach to increase sample sizes while keeping costs fixed^11,12^. Additionally, multiplexing allows for better detection of heterotypic doublets. When samples are genetically distinct, they can be pooled without additional experimental approaches required by antibody- or lipid-based hashing strategies^13,14^. Previous computational approaches developed for droplet-based single-cell RNA-seq experiments leverage genetic variation as a natural barcode in each cell to assign droplets to individual donors^12,15,16^. However, these methods are designed to run on a single modality, and do not leverage both RNA and ATAC reads in the same droplet. Furthermore, these methods typically require prior specification of candidate droplets. Genetic multiplexing offers the advantage to detect empty droplets based on variant calls, which may be superior to detection via expression or accessibility.

An additional challenge encountered with RNA and ATAC single-cell multiome experiments is the potential contamination of exogenous ambient molecules in droplets^14,17–19^. This issue is more prominent with nuclei, especially if isolated from solid or frozen tissues^20^. As current single-cell multiome protocols rely on nuclei isolation^9^, methods to evaluate ambient molecule contamination would be valuable for quality control or covariate correction purposes.

To address these limitations, we developed ambimux, a computational approach to demultiplex single-cell multiome experiments and estimate ambient molecule fractions for each modality. Our method estimates ambient molecule fraction per modality, providing a valuable filtering metric. Additionally, ambimux classifies empty droplets based on genetic data, removing the need for prior filtering. We demonstrate via simulations that ambimux outperforms existing methods for demultiplexing contaminated multiome experiments, recovering a higher number of singlets with greater accuracy. We also show that ambimux accurately estimates ambient fractions in RNA and ATAC modalities in these synthetic data. Finally, we apply ambimux to a single-cell multiome dataset from seven human visceral adipose tissue samples in a single run. Overall, ambimux provides a fast and scalable approach for demultiplexing single-cell multiome experiments and accounting for ambient molecule contamination.

## Results

### Overview of the ambimux model

Ambimux is a likelihood-based method designed for demultiplexing single-cell multiome RNA- and ATAC-seq experiments, while simultaneously accounting for ambient contamination. It offers two novel features that are particularly beneficial for droplet filtering. First, ambimux can classify all generated droplets as empty, singlet, or doublet, eliminating the need for prior barcode filtering. Second, it estimates the fraction of ambient RNA and DNA molecules in each droplet, enhancing assessment and filtering for downstream analyses. Ambimux accomplishes this by using variant-overlapping base calls within individual reads along with sample genotypes without relying on expression or accessibility data.

### Ambimux accurately demultiplexes samples in contaminated simulated multiple data

We hypothesized that explicit modeling of ambient molecules per droplet would lead to improved demultiplexing. To test this, we simulated three multiome datasets with low (mean = 10%), medium (mean = 20%), and high (mean = 30%) ambient fractions (Methods; Fig. 1a). Each simulation consisted of 10,000 nuclei with a 10% doublet rate and eight samples multiplexed uniformly (see methods). We ran ambimux and compared the performance with three other demultiplexing methods: Demuxlet^12^, Vireo^15^, and SouporCell^16^. We ran these methods in each of the RNA and ATAC modalities separately and ran ambimux jointly on both. First, we compared precision across all methods by calculating the percent of classified singlets with a correct donor assignment. Ambimux, Demuxlet, and Vireo had a precision of greater than 99% across each of the three ambient simulations (Supplementary Fig. 1a,b). The precision of SouporCell was somewhat lower, ranging from 92%-98% and decaying slightly with increasing contamination (Supplementary Fig. 1a,b). Overall, these results suggest that demultiplexing under ambient contamination is not highly susceptible to false positives.

**Figure 1:**
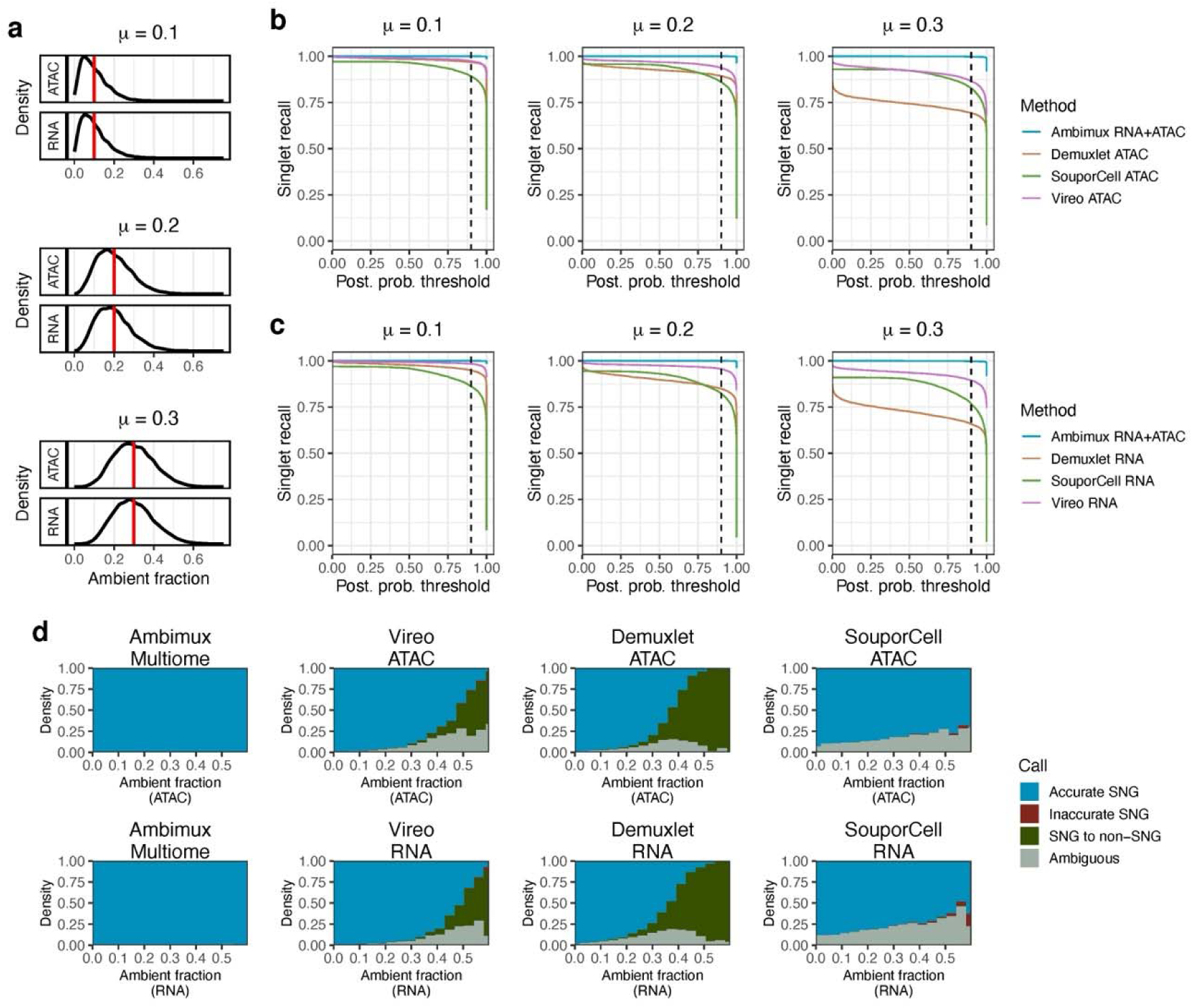
Ambimux recovers more contaminated singlets in a simulated multiome experiment. **a**, Distribution of the proportion of ambient reads in the low (top), medium (middle), and high (bottom) simulated single-cell multiome datasets. The vertical red lines indicate the mean (0.1, 0.2, and 0.3). **b, c,** Recall (sensitivity) of singlet assignments in the three simulated datasets from ambimux, Demuxlet^12^, SouporCell^16^, and Vireo^15^. Ambimux was run on the combined modalities, while all other methods were applied to ATAC (**b**) and RNA (**c**) separately. The dashed vertical line shows the 90% posterior probability threshold. **d,** Relationship between droplet ambient fraction and classification accuracy across methods and modalities. The stacked bars show the proportion of singlets (SNG) classified as accurate (assigned to corrected donor), inaccurate (incorrect donor), non-singlets (assigned as doublet or empty), and ambiguous.

Next, we compared recall (sensitivity) across the methods by calculating how many singlets each method could recover. Of the 9,000 singlets in each of the three contamination datasets, we calculated the percent of these assigned as singlets to the correct donor. We observed that ambimux was able to accurately classify over 99% of the 9,000 singlets in each of the low, medium, and high ambient datasets (Fig. 1b,c). In contrast, all other methods showed a lower recovery of singlets in the higher contamination datasets (Fig. 1b,c). For example, Vireo RNA, the next best method, had a sensitivity of 98.4% in the low contamination data, but only 89.4% in the highly contaminated singlets. Thus, we found that ambient contamination generally decreased the ability to recover singlets when not accounted for.

As both coverage and contamination can likely affect demultiplexing, we sought to identify the contribution of each to the ability to accurately assign singlets. First, we evaluated at what point ambient contamination started to reduce sensitivity. For both Vireo and Demuxlet, sensitivity started to degrade at around 25% ambient contamination, and most singlets were not recovered at around 50% (Fig. 1d). With SouporCell, 20-30% of calls were ambiguous across the entire range of ambient contamination, even at 0% (Fig. 1d). In contrast, ambimux maintained over 99% accuracy in singlet calls even when ambient fractions approached 60% (Fig. 1d). Droplet coverage also affected singlet recovery in single-modality runs. In both Vireo, Demuxlet, and SouporCell, lower coverage led to more ambiguous calls, especially with droplets containing less than 1,000 UMIs or fragments (Supplementary Fig. 1c). In contrast, ambimux was largely unaffected as both modalities contributed reads for demultiplexing (Supplementary Fig. 1c).

### Accurate estimation of droplet-specific ambient contamination

Multiplexing of single-cell experiments with genetically distinct individuals provides a unique opportunity to assess ambient molecule contamination. The distinct “genotype” of the ambient pool allows for probabilistic discrimination between donor and ambient molecules. To leverage this, we incorporated the droplet fraction of ambient molecules as a parameter (see methods). To test the accuracy of these contamination estimates, we combined the three simulated datasets above and compared the estimates with the simulated true values. Overall, we found these estimates to be highly accurate in both modalities (Fig. 2a,b), with a mean absolute error (MAE) of 4.8% and 4.2% in the RNA and ATAC, respectively (Fig. 2a). We further hypothesized that accuracy in droplet ambient estimation would be influenced by coverage. To test this, we fit a local loess regression curve between the droplet absolute error and the number of variant-overlapping reads (informative reads). As expected, we found lower errors in ambient estimates with increasing coverage, with similar parameters in both modalities (Fig. 2c; Supplementary Fig. 2). For example, droplets with 100 informative RNA reads were predicted to have an error of 7.3%, while those with 1,000 informative RNA reads had a predicted error of only 2.7% (Fig. 2c).

**Figure 2:**
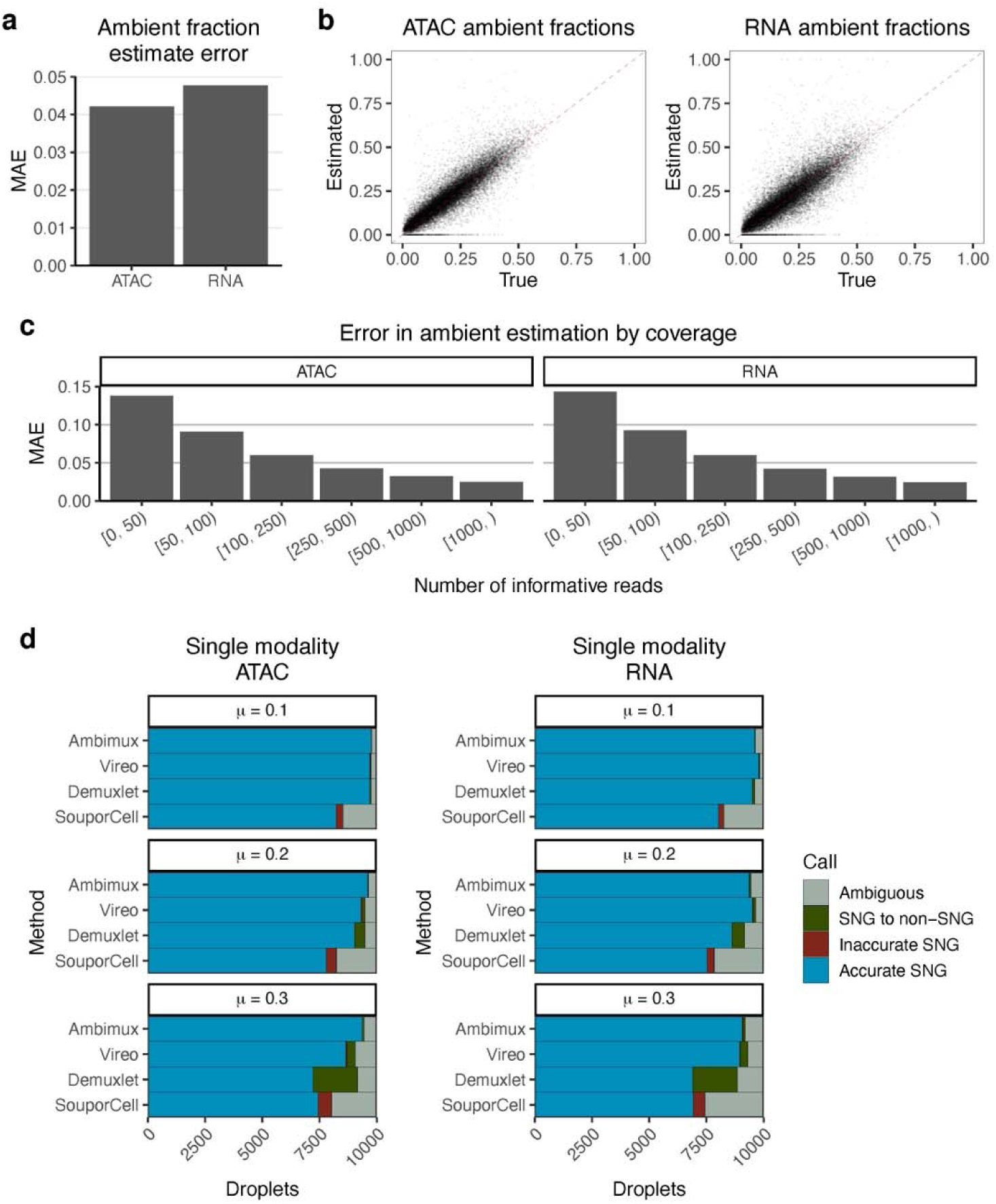
Accurate estimation of ambient proportions in simulated multiome datasets. **a**, Mean absolute error (MAE) of ATAC and RNA ambient proportion estimates by ambimux after combining results from the three simulated datasets of low (mean 0.1), medium (mean 0.2), and high (mean 0.3) contamination. **b,** Correlation between true and estimated ambient fractions for the ATAC (left) and RNA (right) modalities in the combined results. **c,** MAE of ambient fraction estimates grouped by coverage, demonstrating improved estimation accuracy with increasing coverage. **d,** Comparison between ambimux and competing methods in singlet (SNG) classification when run on either ATAC (left) and RNA (right) data separately. Results are grouped by ambient dataset. The bar plots show the proportion of ground truth singlets assigned as correct singlets, incorrect singlets, incorrect doublets/empty, and ambiguous.

We also compared our background fraction estimates with that of CellBender^18^, a deep generative model for ambient reads removal in single-cell RNA-seq using count data. As our simulations generated a distinct background distribution, we reasoned that CellBender could be applied to our three synthetic contaminated datasets. In addition to RNA, we tested ATAC counts, although we note that the authors never evaluated CellBender in this modality. We found that CellBender failed to properly estimate background fractions (Supplementary Fig. 3). After combining the results from the three datasets, the mean absolute errors were 19.0% and 18.4% in the RNA and ATAC modalities, respectively (Supplementary Fig. 3a). Upon further inspection, we found that CellBender underestimated the ambient fraction on average, and the MAE increased with higher background reads (Supplementary Fig. 3b). Importantly, background estimates from CellBender were uncorrelated with the simulated ambient fractions for each of the 3 ambient datasets (Pearson R < 0.01) (Supplementary Fig. 3c). Overall, these comparisons highlight how modeling genotypes can improve estimation of ambient read fractions when compared to read counts alone.

### Ambimux improves demultiplexing of contaminated droplets in single modalities

While developed for single-cell multiome data, ambimux can easily run on a single modality. We tested ambimux on the RNA and ATAC modalities separately, using the three simulated datasets described above (low, medium, and high contamination). This allowed us to assess the benefits of ambient modeling independently of coverage, as the RNA+ATAC approach doubled the number of reads on average compared to the single-modality approach. We again compared ambimux with Demuxlet^12^, Vireo^15^, and SouporCell^16^ and assessed sensitivity in accurately recovering the 9,000 singlets in each dataset (Fig. 2d). In the RNA modality, Vireo accurately recovered the most singlets in the low (98.4%) and medium (95.5%) contamination datasets, while ambimux accurately recovered 97.3% and 95.2%, respectively (Fig. 2d). In the high contamination dataset, ambimux performed best, accurately assigning 92.5% of singlets, whereas Vireo recovered 89.4% (Fig. 2d). Ambimux also outperformed all methods in the ATAC modality, recovering 98.3%, 97.3%, and 95.2% of the 9,000 singlets in the low, medium, and high ambient datasets, respectively (Fig. 2d). These results show that ambimux is robust to ambient contamination even when run on either RNA or ATAC modalities individually, although joint RNA+ATAC calling performed best (recall > 99%) in all three simulated datasets.

### Ambimux maintains accuracy across pooling strategies

An important question in the design of multiplexed experiments is how many samples to pool. Often, this will involve balancing depth vs. breadth with budgetary constraints^21^. We sought to test the performance of ambimux across a wide range of pooling numbers. We simulated synthetic data sets consisting of 2, 4, 8, 16, 32, and 64 pooled samples. As before, each pooling experiment contained 9,000 singlets and 1,000 doublets. Droplet background fractions were sampled from an equal mixture of low, medium, and high ambient distributions (Methods). We observed that ambimux maintained excellent performance across all pooling numbers, with precisions and recalls greater than 99.9% and 99.8%, respectively (Fig. 3a). We also found that ambimux could accurately estimate ambient fractions with mean absolute errors of 4.6-6.1% for the RNA and 4.1-5.3% for the ATAC (Fig. 3b). Interestingly, the accuracy of background fraction estimates increased with higher pooling numbers (Fig. 3b).

**Figure 3:**
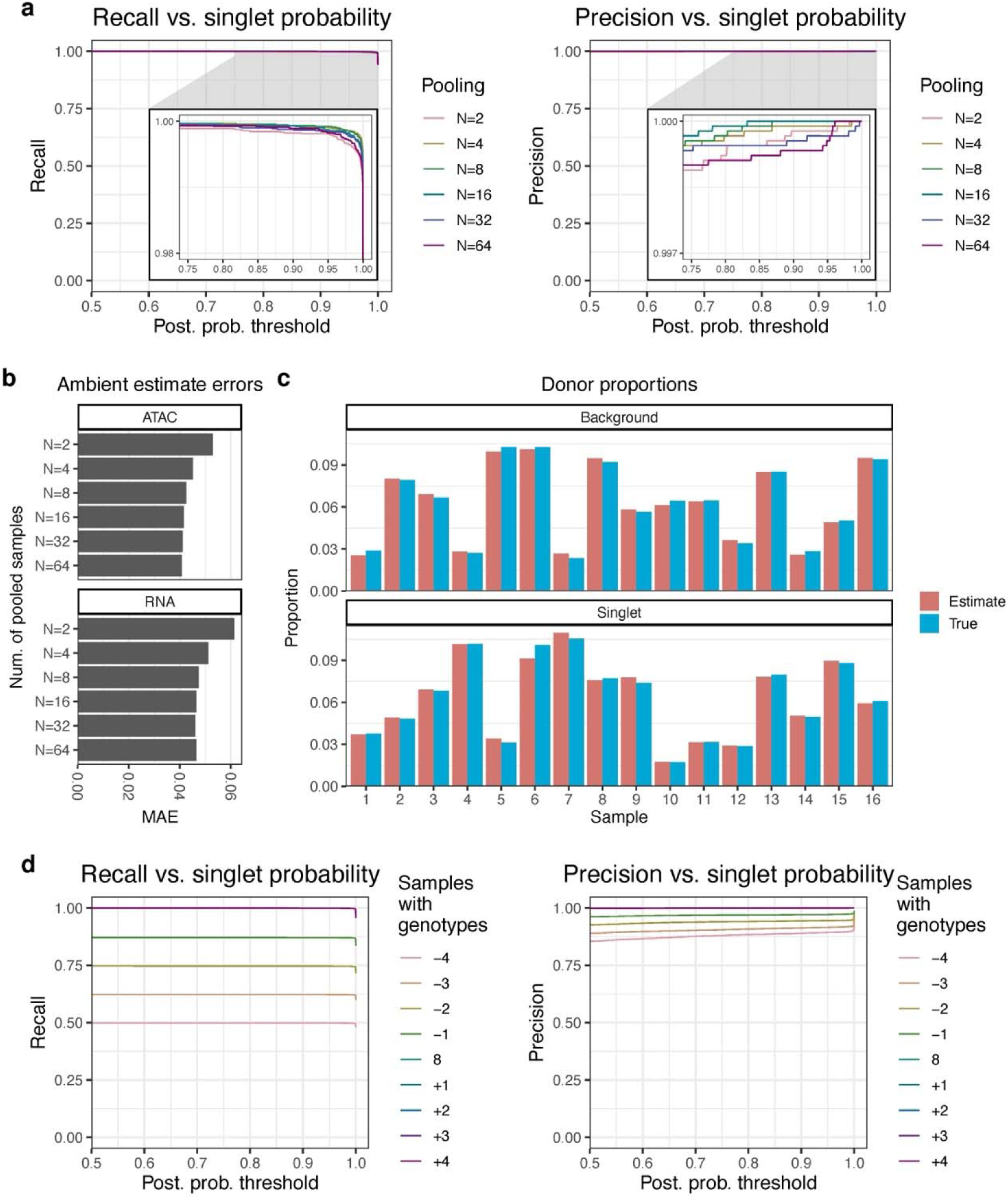
Ambimux is robust to variations in pooling numbers and sample dropout. **a**, Recall (left) and precision (right) curves for singlet assignment when pooling 2, 4, 8, 16, 32, and 64 simulated genetically distinct donors from 1000 Genomes^36^. The x-axis shows the singlet posterior probability threshold while colors indicate pooling number. **b,** The mean absolute error (MAE) for ATAC (top) and RNA (bottom) of the ambimux ambient fraction estimates based on the number of pooled samples (y-axis). **c,** Comparison of ground truth and estimated donor proportions in the background (top) and nuclei (bottom) pool in a simulated pool of 16 donors. The data were generated such that the proportion of the 16 donors in the background pool was independent of that of the singlets. **d,** Recall (left) and precision (right) curves for ambimux demultiplexing with simulated donor experimental dropout (adding genotype samples) and missing genotypes (removing genotype samples). Ambimux was run on the same simulated pool of eight donors but varying the genotype samples used for demultiplexing, either removing donors (negative numbers) or adding donor genotypes (positive numbers).

To properly model ambient molecules in a droplet, ambimux requires the allele frequencies of the ambient pool. While donor genotypes are given, the ambient pool “genotype” must be estimated. To do so, we model the ambient allele frequency of a SNP as the average of the donor genotypes weighted by estimated background donor proportions (see methods). We thus asked whether our model can properly estimate these fractions of donors in a pool. In a simulation of 16 samples with a 10-fold variation in abundance, we found that the estimated donor proportions accurately reflected the true simulated values in both the background pool and singlet droplets (Fig. 3c). This allowed for accurate estimation of ambient fractions with an MAE of 0.041 and 0.046 for the ATAC and RNA, respectively (Supplementary Fig. 4a). Additionally, ambimux maintained high precision (> 99.9%) and sensitivity (99.9%) in this simulation (Supplementary Fig. 4b,c). Our results on these simulated data show that ambimux is robust to variations in sample yield, both in the ambient and singlet droplet pools.

Next, we asked whether ambimux could accurately demultiplex datasets in which genotypes were missing or samples dropped out. We tested this by simulating a multiome dataset of eight donors and demultiplexing after removing donor genotypes or adding new donor genotypes in addition to the 8. We performed a sweep by removing 1, 2, 3, and 4 sample genotypes, simulating cases of missing genotypes, and adding 1, 2, 3, and 4 sample genotypes, simulating cases with experimental dropout. We found that the addition of sample genotypes had minimal effects on recall and precision (Fig. 3d), suggesting ambimux can accurately detect and assign singlets after dropout of samples in a pool. When genotypes of pooled donors were missing, recall of the genotyped donors was largely unaffected. For example, the recall from demultiplexing 6 out of 8 samples was 0.747, near the maximum of 0.75 (Fig. 3d). However, missing genotype samples detrimentally affected the ability to accurately assign singlets to donors (Fig. 3d). The precision was 0.890 when only 4 of the 8 genotypes were present, while the precision was 0.999 with the full data (Fig. 3d). Missing or added genotypes also slightly worsened the accuracy of ambient estimates. While demultiplexing with the eight pooled samples resulted in an MAE of 0.043 and 0.047 for the ATAC and RNA respectively, demultiplexing with 4 missing genotypes resulted in MAEs of 0.088 and 0.091, for example (Supplementary Fig. 5a). With missing genotypes, we found that there were singlets with higher, overestimated ambient fractions above 0.5 (Supplementary Fig. 5b), likely originating from ungenotyped samples assigned to an incorrect donor and fit with a high ambient fraction.

### Ambient contamination reduces power to detect differential abundance

Many single-cell analyses involve detecting feature-level differences (such as gene expression or ATAC accessibility) between conditions^21^, including cell-type-specific marker identification and disease association^22^. We investigated the extent to which ambient contamination can affect differential abundance (DA) analysis of peaks or genes. To test this, we generated three synthetic datasets with low (mean 10%), medium (mean 20%), and high (mean 30%) background contamination levels. Each dataset comprised eight pooled individuals, with four containing a disease cell subtype. We simulated various log fold-changes for DA features in the disease cell-type, ranging from 0.1 to 2.0 for 1,000 genes and 1,000 peaks (Fig. 4a). DA of the 1,000 features in each modality showed that ambient contamination decreased the power to detect differential abundance (Fig. 4b). Specifically, in the RNA modality, we detected 334, 286, and 234 DA genes in the low, medium, and high ambient datasets, respectively (Fig. 4b). The ATAC modality showed a similar trend, with 27, 20, and 10 DA peaks detected (Fig. 4b). As expected, features with high fold-changes were most detectable, and estimated log fold-changes correlated with the true log fold-changes for each dataset (Supplementary Fig. 6).

**Figure 4:**
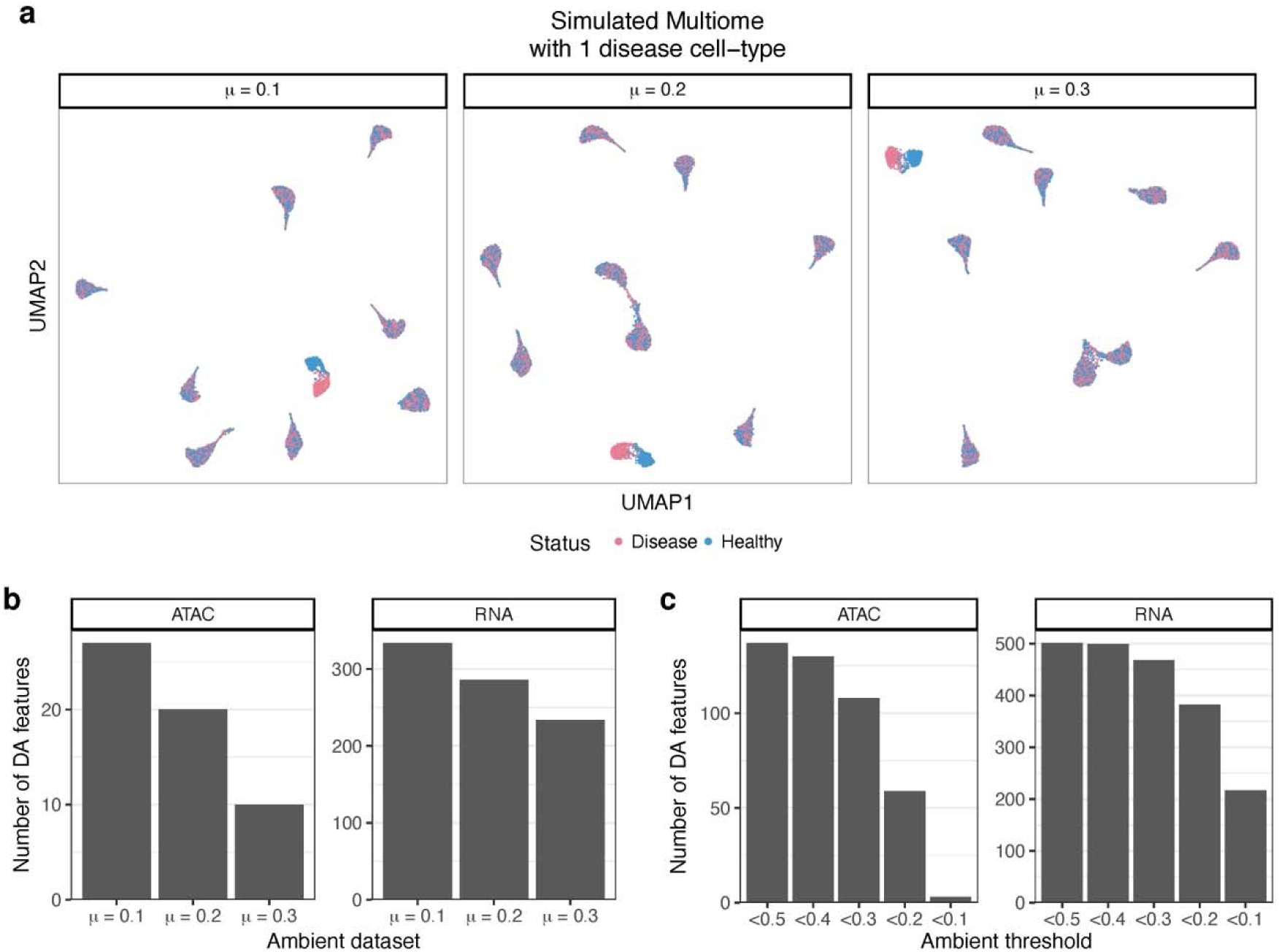
Ambient contamination reduces power to detect differentially abundant peaks or genes. **a**, UMAP visualization of three simulated datasets with ambient fraction means of 10% (left), 20**%** (middle) and 30% (right). Each dataset was simulated from the same 9 cell-types. We artificially added a disease condition for four of the eight donors, where we introduced varying log fold-changes for 1,000 peaks and 1,000 genes in one cell-type. **b,** Number of significant differentially abundant (DA) features (Bonferroni < 0.05) in the disease cell-type between conditions for each of the three datasets. This shows ambient contamination decreases power to detect significant DA features. **c,** Number of significant DA features (Bonferroni < 0.05) as in (**b**), but after filtering out droplets by various ambient fraction thresholds in the combined data. The three ambient datasets were merged, and DA feature testing was performed after filtering by the ambient fraction threshold indicated in the x-axis.

After confirming that higher background levels led to decreased power to detect DA, we sought to understand how filtering out contaminated droplets could affect DA results. To investigate this, we combined the three synthetic datasets above and tested DA after removing droplets above various ambient fraction thresholds. We used a range of 0.1 to 0.5 for ambient fraction removal. Filtering out the fewest droplets at a threshold of 0.5 resulted in the highest number of DA genes, with 137 DA peaks and 501 DA genes in the ATAC and RNA modalities, respectively (Fig. 4c). In contrast, removing singlets with an ambient fraction above 0.1 resulted in a much lower number of significant features, with 3 DA peaks and 217 DA genes (Fig. 4c). These observations are likely due to the fact that ambient contamination and healthy-disease states were simulated independently of each other, such that contamination was not a confounder for DA testing. Overall, these results suggest that ambient contamination reduces power to detect differential feature abundance, and further filtering of features also reduces power.

### Characterizing ambient contamination and filtering in a single-cell multiome dataset from visceral adipose tissue

Encouraged by our results on synthetic data, we next investigated the performance of our method on observed data. We ran the 10x single-cell Multiome assay on seven pooled visceral adipose tissue (VAT) samples from the KOBS cohort^23,24^. After mapping with CellRanger Arc v2.0.2^25^, ambimux identified 4,986 singlets and 512 doublets (9.3%). Several test droplets with low coverage were classified as empty, and droplets with a low coverage in only one modality could be assigned to a singlet with a higher coverage in the other modality (Fig. 5a).

**Fig. 5:**
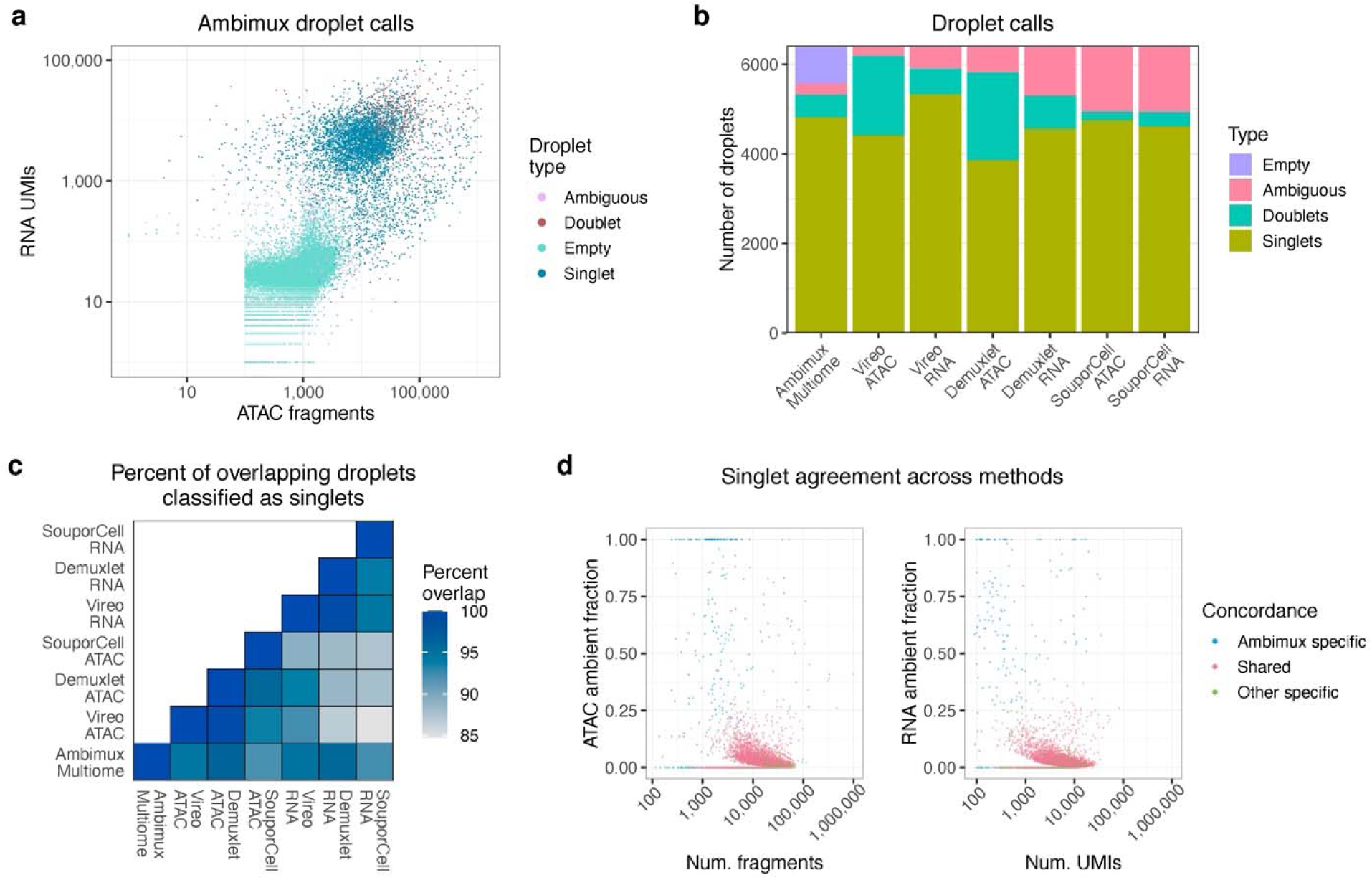
Ambimux demultiplexing of seven visceral adipose tissue samples. **a**, Droplet classification by ambimux run on multiome data generated from seven pooled visceral adipose tissue samples. Low coverage droplets tended to be classified as empty, while higher count droplets tended to be classified as doublets. **b,** Classification of 6,414 candidate droplets across methods. Ambimux was run on RNA+ATAC, while Vireo, Demuxlet, and SouporCell were run on either ATAC or RNA. Note that only ambimux is able to assign empty droplets. **c,** Concordance of singlet classification between methods. The overlap is defined as the number of droplets classified as singlets by both methods divided by the minimum number of singlets called by either method. **d,** Scatterplot showing singlet agreement between ambimux and the three methods from above. Singlets are plotted against estimated ambient fraction and read coverage for ATAC (left) and RNA (right) and colored by whether the singlet was called by ambimux only (blue), called by both ambimux and at least one other method (red), or called by at least one other method only (green).

Next, we compared ambimux droplet calls with those from demuxlet, Vireo, and SouporCell. After restricting analysis to 6,414 candidate droplets with at least 100 UMIs and fragments, ambimux identified 4,824 singlets, while other methods classified a range of 3,857 to 5,334 singlets (Fig. 5b). On average, 92.4% of droplets called as singlets in one method were called as singlets in another method (Fig. 5c). Ambimux singlets overlapped with at least 94.2% of singlets called in the other approaches and showed slightly higher overlap with ATAC-based demultiplexing (Fig. 5c). Among the intersecting singlets, methods had high concordance (mean = 98.5%) in their sample assignments (Supplementary Fig. 7a). We then compared singlets that were unique to ambimux when compared to other methods applied to either ATAC or RNA. The 270 (ATAC) and 183 (RNA) droplets specific to ambimux tended to have higher contamination scores or lower coverage (Fig. 5d). On the other hand, singlets specific to the other methods tended to be called as doublets by ambimux (Supplementary Fig. 7b).

We considered that lysed or broken nuclei could release randomly fragmented DNA with loss of chromatin structure and thus be accessible to the Tn5 transposase. This could lead to extranuclear DNA preferentially originating from inaccessible regions that make up most of the genome. We tested this in the VAT multiome dataset by using either fragments within peaks (intra-peak) or between peaks (inter-peak) for demultiplexing. In line with our hypothesis, the mean ATAC ambient fraction was higher from the inter-peak fragments (22.4%) compared to those from the intra-peak fragments (6.3%) (paired Wilcoxon p < 2.2x10^-16^) (Fig. 6a). While higher than the intra-peak ambient fractions, the inter-peak proportion (22.4%) is lower than the fraction of the genome that these regions occupy (97.3%). Notably, the intra-peak contamination estimates were within the same range as the RNA contamination estimates (mean 7.4%) (Fig. 6b). Due to these results, and the fact that downstream analyses rely on peak counts, we used only intra-peak molecules for demultiplexing in this VAT dataset.

**Figure 6:**
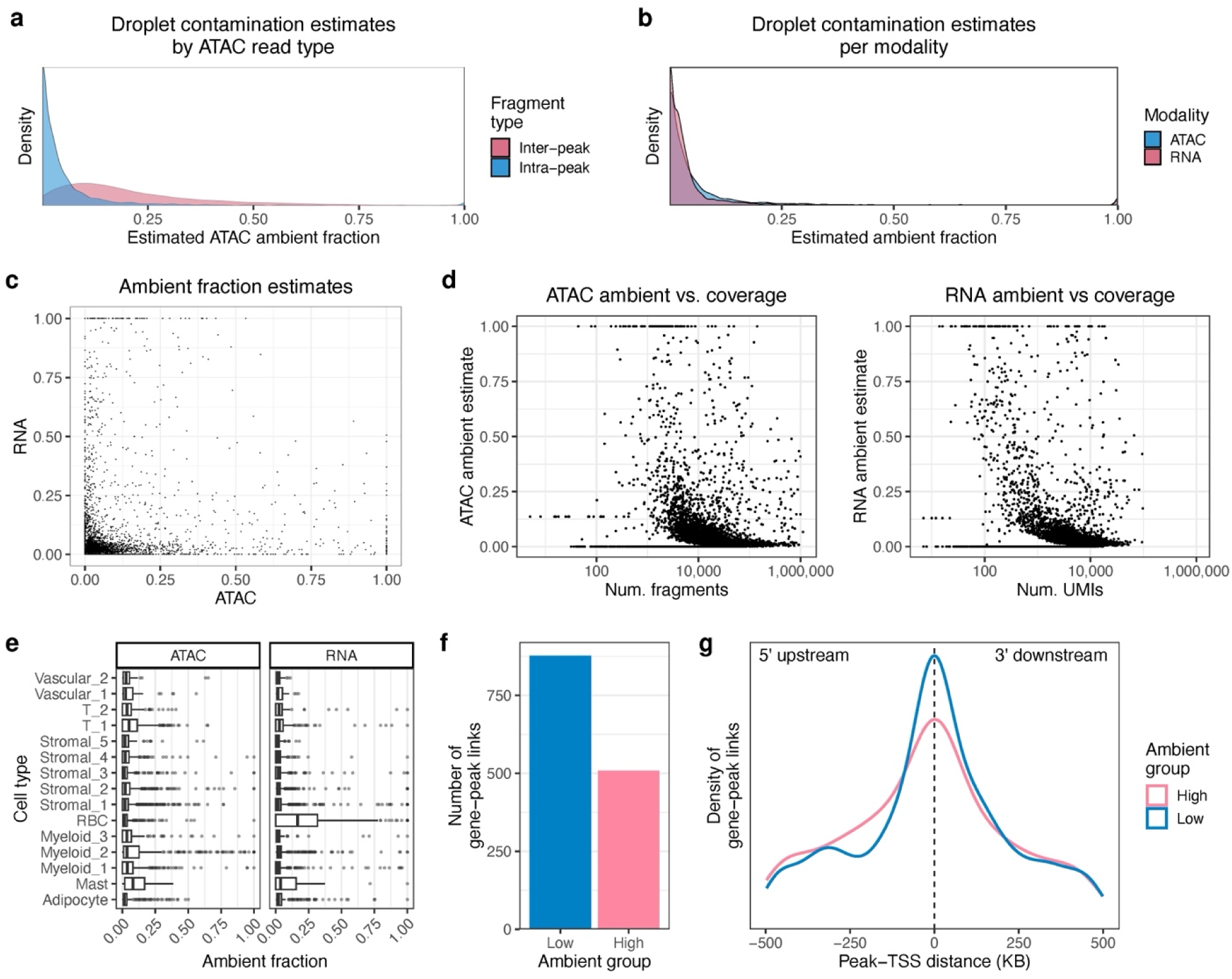
Estimation of ambient fractions in pooled multiome of seven visceral adipose tissue donors. **a**, Distribution of ambient fraction estimates from inter-vs. intra-peak reads. Inter-peak read ambient fraction is estimated from reads originating from outside peaks, while intra-peak read ambient fraction is estimated from reads inside peaks. **b, c,** Relationship between ATAC and RNA ambient fraction estimates in singlets. The density plot shows a similar distribution of ambient fractions for both modalities (**b**), while the scatterplot shows a low correlation between the two (**c**). **d,** Correlation between read coverage and ambimux ambient fraction estimates in singlets for ATAC (left) and RNA (right). **e,** Distribution of ambimux ambient fraction estimates per cell-type in the ATAC and RNA modalities. **f, g,** Number of significant gene-peak links (FDR-corrected p < 0.05) in low- and high-ambient droplets (**f**), and the distance between the peak and transcription start site (TSS) for these gene-peak links (**g**). Low- and high-ambient droplets were defined as those with less than 5% or more than 5% estimated ambient percent in both modalities, respectively.

Next, we investigated whether RNA and ATAC ambient fractions were correlated with common summary statistics in visceral adipose tissue singlets. Notably, the ambient fractions between modalities were weakly correlated (Fig. 6c), with a Spearman coefficient of 0.08 (p = 4.9 x 10^-8^), suggesting that background contamination of RNA and ATAC molecules occurs independently. We also found that lower coverage droplets tended to have a somewhat higher amount of contamination in both modalities (Fig. 6d), with Spearman correlations of -0.11 and -0.13 for the ATAC and RNA, respectively. This is consistent with a model of uniform ambient molecule abundance that would result in low coverage droplets containing a higher proportion of background. Lastly, we found that the percent of mitochondrial reads, a typical filtering and QC metric^8,18^, showed only slightly positive correlations with ambient contamination, with ATAC and RNA Spearman correlations of 0.10 and 0.21, respectively (Supplementary Fig. 8a).

Finally, we were interested in seeing how ambient fractions could affect downstream analysis. To do so, we performed gene-to-peak link analysis on clean and contaminated visceral adipose tissue singlets. First, we clustered droplets and assigned them to 15 cell-types, including vascular, stromal, myeloid, and adipocyte cells (Supplementary Fig. 8b). We found that ambient contamination varied across cell-types (Fig. 6e), although this was largely correlated with cell-type coverage (Supplementary Fig. 8c). We then split the nuclei into equally sized low ambient (< 5% in both RNA and ATAC) and high ambient (> 5% in both RNA and ATAC) droplets (N = 167), controlling for read depth and cell-type (Supplementary Fig. 9). We found 877 and 508 significant links (FDR-adjusted p < 0.05) in the low and high groups respectively (Fig. 6f). Notably, the associated peaks in the clean droplets tended to be closer to the transcription start site (TSS) than those of the contaminated droplets (Fig. 6g). Overall, these results suggest that ambient contamination can degrade the power to detect significant gene-peak links and lead to potentially more spurious associations.

## Discussion

We developed ambimux for integrated demultiplexing of single-cell multiome experiments under ambient contamination. We show that our method outperforms existing approaches in sensitivity and specificity. Using synthetic data, we were able to show how our approach correctly handles experiments with high ambient contamination and identify points where other methods break. As part of the model, ambimux also outputs modality-specific ambient fraction estimates per droplet. Our results on simulations show that ambimux can accurately estimate droplet contamination with mean absolute errors as low as 2.7% for higher coverage droplets. We also show that ambimux is capable of demultiplexing a wide range of pooling numbers, from just 2 donors to 64. Importantly, ambimux is robust to donor proportion variation, and is robust to sample dropouts and missing genotypes. Finally, we applied ambimux to a real dataset of seven pooled VAT samples and showed comparable results to competing methods.

While highly optimized protocols and cell line samples can yield higher quality data, intact cell or nuclei isolation from tissue remains challenging in many cases^26^. Furthermore, generating cDNA single-cell libraries from fresh biopsies can present logistical challenges, particularly in scaling to larger sample sizes. Therefore, frozen tissues might be the only practical approach. However, frozen tissues are particularly subject to lysis and can lead to increased ambient molecule concentrations^20,26^. As single-cell RNA, ATAC, and multiome methods are applied to frozen tissues, ambient fraction estimates from genetic data can provide a high confidence method to assess the quality of the experiment, filter individual cells, and correct for potential confounding. We have shown here that ambimux can accurately estimate these contamination fractions across a wide range of experimental setups.

We hypothesize that ambimux will help enable population-scale single-cell multiome studies in tissues. By providing a robust method to demultiplex samples and estimate background contamination, ambimux allows researchers to increase pooling and sample sizes for costly multiome experiments. This approach can help to better identify subtle regulatory mechanisms across populations^21,27^. Moreover, by providing grounded contamination estimates for use as a covariate or filtering threshold, ambimux may help improve the accuracy of downstream analyses, such as peak-to-gene links. Improved analyses will help advance our understanding of cell-type-specific gene regulation and offer new insights into developmental processes, disease mechanisms, and tissue heterogeneity^28,29^.

Despite its advantages, ambimux has some limitations. First, the method relies on genetic variation between samples, which may limit its applicability in cases where samples are genetically identical, such as cell lines, or when working with model organisms with limited genetic diversity. Additionally, we observed a drop in ambient estimation accuracy in low coverage droplets, as sparsity of informative reads can lead to high variance in parameter inference. Finally, ambimux was designed for multiplexed experiments in which all donors have genotype data available. In practice, it may be challenging to obtain this for every donor. While we observed that ambimux is robust to some missing genotypes, explicit modeling of this may be preferable.

Looking ahead, several promising avenues for further development of ambimux can be identified. A key area is the extension of the model for ambient molecule correction. Ambimux already models ambient contamination at the read level internally. By leveraging this modeling approach, functionality could be developed to correct and remove ambient molecules from read counts, similar to the approach used by CellBender^18^. This would provide a more accurate representation of true cellular content, potentially improving downstream analyses such as differential feature abundance. As alluded to earlier, an extension of a genotype-free or missing-genotype framework would further expand the utility of ambimux in these cases.

In conclusion, ambimux represents a significant addition to the single-cell field. By offering multimodal ambient contamination estimates, ambimux addresses common challenges faced with large-scale multiome studies in tissues. Its ability to handle various experimental designs and contamination levels, coupled with its potential to facilitate larger-scale studies, positions ambimux as a valuable tool for single-cell studies of gene regulation.

## Methods

### Model Description

The foundation of our approach is to model the observed variant-overlapping base calls in reads using a Bernoulli distribution. Our Bernoulli modeling approach is inspired by the work of Kang *et al*^12,30^, and we extend this by incorporating parameters for empty droplets and ambient reads. We take a likelihood-based approach and use posterior probabilities to assign donors to droplets. Since the model is the same for both RNA and ATAC modalities, we use the term ‘molecule’ to refer to either a deduplicated ATAC-seq fragment or a deduplicated RNA-seq unique molecular index (UMI). We first define the components of the probability model and then describe the estimation procedure.

In a multiome experiment, let *D* be the number of droplets containing at least one molecule. A droplet *d* can have either 0, 1, or 2 nuclei encapsulated, denoted by the latent variable *H_d_*. We assume that when multiple nuclei are present in a droplet, their combined genotype resembles that of the ambient pool, so we do not consider cases with 3 or more nuclei. This assumption simplifies the estimation process by avoiding the complexity of modeling higher order combinations.

Although the number of cells or nuclei encapsulated in a droplet roughly follows a Poisson distribution, nuclei can adhere to each other, resulting in higher rates of doublets^31^. To account for this, we model *H_d_* using a categorical distribution parameterized by *λ*.

Let *N* be the number of samples pooled in an experiment. The composition of donors in a droplet, denoted by *S_d_*, is conditional on the droplet type *H_d_*. There are three possible droplet types: empty droplets containing no donors, singlets containing one donor, and doublets containing two donors. The experiment-wide probability of obtaining a singlet sample *{i}* or doublet samples *{i, j}* is modeled using a Categorical distribution parameterized by π*_c_*, which is a vector representing the proportions of each donor in the cells/nuclei.

For a singlet, the probability of being assigned to donor *i* is given by *π_ci_*. For a doublet, the probability of being assigned to donors *i* and *j* is given by the product *π_ci_π_cj_*, assuming that the probability of each donor in a doublet is independent of the other donor. To improve efficiency, self-doublets are ignored, and only unique doublet possibilities are modeled.

Assume a droplet *d* contains *M_d_* molecules. For a singlet or doublet, each molecule *m* can originate from a sample *s_i_*(*i* = 1, …, *N*) or from the ambient pool *s_0_*. We introduce the latent random variable *T_dm_* to indicate the origin of each molecule, which can be either a single sample or the ambient pool. The probability of a molecule originating from the ambient pool is modeled as a Bernoulli distribution with parameter *α_dhs_*, where *h* and *s* denote the specific *H_d_* (number of cells) and *S_d_*(sample identity) for the droplet. If the droplet is empty (i.e., *H_d_* = 0), then *α_dhs_* = 1. For a doublet, we assume that each donor sample has an equal probability of contributing a molecule. Note that we specify a droplet contamination parameter *α_dhs_* specific to each combination of *H_d_* and *S_d_*. Thus, for a clean singlet, the parameter estimate will be close to 0 for its assigned donor but should be much higher for an incorrect donor assignment.

We model the probability of observing a base call at variant sites within a read. Let *B_dmv_* represent the observed base call at variant site *v* in molecule *m* of droplet *d*. Base calls *b* can be either 0 (reference allele) or 1 (alternate allele). Our method focuses on bi-allelic SNPs, disregarding other types of genetic variants.

The observed base depends on whether a sequencing error occurred, represented by the latent variable *E_dmv_*. Sequencing errors follow a Bernoulli distribution with probability *τ_dmv_*, derived from the Phred quality score of the base call.

In the absence of an error (*E_dmv_* = 0), the observed base call follows a Bernoulli distribution parameterized by *γ_iv_*, which is the alternate allele frequency of variant *v* for origin *i* (donor or ambient pool). Donor allele frequencies are provided, while ambient allele frequencies are estimated from the data. In the event of a sequencing error, we assume equal probability for observing any base.

Taken together, the probability of a droplet is given by

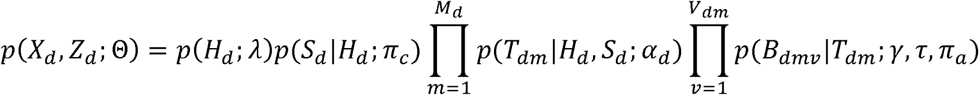

where *X_d_* gives the observed data, *Z_d_* gives the latent variables *H_d_*, *S_d_*, *T_d1_*, …, *T_dm_*, and *Θ* gives the parameters *λ*, *π_c_*, *α*, *π_a_*, *γ*, and *τ*.

### Parameter estimation and model fitting

The purpose of our method is threefold: 1) identify empty, singlet, and doublet droplets, 2) assign droplets to donors, and 3) estimate ambient fractions in each droplet. We achieve this using a combination of gradient-based methods and expectation maximization (EM) ^32^. The output of interest consists of the posterior probabilities of *H_d_*and *S_d_*, and the modality-specific droplet ambient estimates *α_d_*. While all droplets are input to the model, we only estimate parameters for droplets with at least U = 100 molecules in either modality, treating all others as empty, consistent with previous works^20,33^. For single-modality data, the other modality is ignored.

Accurate demultiplexing and ambient estimation relies on the alternate allele frequencies of ambient molecules. Rather than directly calculating empirical allele frequencies from ambient molecules, we model these frequencies as a function of the donor fractions in the ambient pool and their allele frequencies. This approach regularizes estimates for variants with low coverage and reduces noise. We set the ambient “genotypes” as a weighted sum of donor genotypes: 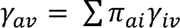 where *γ_iv_* are the given donor genotypes and *π_ai_* are the donor proportions in the ambient pool. We estimate *π_a_* by maximizing the log likelihood of the data using gradient ascent, projecting the updated parameter onto the simplex^34^ to ensure non-negative proportions sum to 1 (Supplementary Note 1). For efficiency, we calculate *π_a_* using only fixed empty droplets and keep this estimate constant.

To estimate droplet ambient parameters *α_dhs_* for all test droplets across *H_d_* and *S_d_*, we use Newton-Raphson optimization for each droplet independently. We maximize the log likelihood after marginalizing over *T_dm_*, adding a prior to avoid collapsing the likelihood (Supplementary Note 1).

With updated values for *α_dhs_*, we estimate *λ* and *π_c_* by maximizing the expected log likelihood, where the expectation is taken with respect to the posterior of the latent variables. Each iteration involves calculating the expected log likelihood and maximizing it with respect to *λ* and *π_c_*. The derivations of the update equations are provided in Supplementary Note 1.

Prior to estimation, we initialize *λ_0_* as the proportion of fixed empty droplets, while *λ_1_* and *λ_2_* are set to 0.9 and 0.1 times the proportion of test droplets, respectively. The values of *π_c_* and *π_a_* are initialized uniformly.

The complete estimation procedure can be summarized as follows:

1. Initialize all parameters and estimate a fixed value for ambient sample fractions *π_a_* from the empty droplets.
2. Iterate until convergence:

a. Estimate ambient fractions *α_dhs_* using Newton-Raphson.
b. Estimate *λ* and *π_c_* by maximizing the expected log likelihood.
c. Set the prior parameter *β* to the weighted average of *α_dhs_* across singlets.
3. Terminate iterations when the mean absolute difference in the parameter estimates is less than *ε* = 10^-6^.

### Simulation of ambient-contaminated single-cell multiome experiments

While methods exist for simulating single-cell experiments^35^, to our knowledge, none can explicitly simulate multiplexed samples with genotypes and controlled ambient contamination. To benchmark ambimux, we developed a simulation method called ambisim^19^. This single-cell multiome simulator generates droplets from a pool of donors, sampling variant alleles during read generation.

Ambisim generates multiome droplets from a set of barcodes with information on donor, cell type, and read types (donor vs. ambient). It produces two sets of droplets: empty and cellular. For empty droplets, the number of reads in both modalities is drawn from a negative binomial distribution (mean = 2, size = 0.1). For cellular droplets, reads are similarly drawn but with mean = 5,000 and size = 1.5, constrained between 200 and 100,000 reads.

In cellular droplets, reads are divided into donor and ambient. The fraction of ambient reads is sampled from a beta distribution, with parameters varying by experiment: low ambient (shape 2,18; mean = 0.1), medium (shape 4,16; mean = 0.2), and high (shape 6,14; mean = 0.3). When evaluating the accuracy of the ambient estimates, we pool the low, medium, and high datasets, unless otherwise stated.

For singlets and doublets, one or two cell types are sampled from a uniform categorical distribution. Genes or peaks are then sampled based on a Multinomial distribution of the randomly drawn cell type(s) profiles.

In the gene expression modality, genes are sampled from the cell type’s expression profile. For ambient reads, an average profile weighted by cell type frequency is used. An isoform is randomly selected with uniform probabilities, then classified as spliced or unspliced (probability 0.4 for cellular, 0.6 for ambient). This reflects the fact that nuclear RNA will be enriched for unspliced mRNA relative to cytoplasmic or total cell RNA. The location of the single-end read sequence is sampled uniformly within the cDNA sequence of the spliced or unspliced isoform.

The ATAC modality employs a similar procedure but accounts for inter-peak reads. Peaks are sampled using a Multinomial distribution with cell type accessibility probabilities. The ambient peak distribution combines cell type distributions weighted by their frequency. For inter-peak reads, its genomic location is sampled randomly from the genome. Whether a read falls inside or outside a peak depends on whether the droplet is empty. For an empty droplet, we sample a read coming from a peak using a Beta distribution with shape parameters (1, 9). For non-empty droplets, we sample a read coming from a peak using a Beta distribution with shape parameters (4, 6). To generate the paired-end read, we sample an insert size from 0 to 150 uniformly.

For both ATAC-seq and RNA-seq, the read sequences are further modified as follows. First, all variants overlapping the read alignment are selected. For each variant in a cellular read, an allele is sampled from a Bernoulli using the allele frequency of the read’s donor and the base at the read site is replaced with this allele. The procedure is similar for an ambient read, where the ambient allele frequency is used instead of the donor’s allele frequency. To obtain the final read sequence we sample a sequencing error with a probability of 0.01 for each base pair. Given an error, the base is replaced randomly with any of the four nucleotides.

For all simulations, we set the cell types and gene and peak probability distributions from the single-cell multiome visceral adipose tissue data set (see below). Briefly, the top 3,000 barcodes ranked by gene expression UMIs were clustered using Seurat v4.3.0^8^ with a resolution of 0.5 and otherwise default parameters. This resulted in nine cell types, and the proportion, gene expression, and peak probabilities for each were calculated empirically. Donors and genotypes were simulated from 1,000 Genomes^36^. We randomly selected unrelated European donors for all simulated data. For each dataset and demultiplexing run, we kept biallelic SNPs and removed SNPs monomorphic in the pooled donors. This set of SNPs was used for both generating the fastq files and demultiplexing donors.

All simulated fastqs were aligned with CellRanger Arc v2.0.2^25^ using GRCh38 and GENCODE v41^37^, the same references from which the data were generated. Additionally, we provided the peaks BED file to the ‘--peaks’ option to ensure that all simulations generated the same peak count data.

### Differential abundance analysis of RNA and ATAC features

To evaluate the effect of ambient contamination on differential expression and differential accessibility, termed differential abundance (DA) here, we slightly modified the data generation to include a disease cell subtype. We took the adipocyte cell-type and modified the multinomial probabilities. We generated log2 fold-changes ranging from -2 to 2 for 1,000 randomly selected features with an expression probability above 1 x 10^-5^. For four individuals we replaced the adipocyte cell-type with the disease cell-type. The three datasets were then generated as described above with low, medium, and high ambient contamination.

To detect DA features, we first demultiplexed the aligned data with ambimux and extracted the assigned singlets. The counts from CellRanger Arc were preprocessed using Signac^38^, where the RNA counts were normalized to sum to 1,000 and log transformed. Finally, we performed a standard Wilcoxon test using the built-in “FindMarkers” function in Seurat^8^ between the healthy and disease cell-type. DA testing was performed only for the 1,000 disease features in each modality. For evaluating the effect of filtering droplets by ambient contamination, we ran FindMarkers using singlets for which the estimated ambient fraction of that modality was below the threshold. We defined significant DA features using a Bonferroni-corrected p-value threshold of 0.05.

### Single nucleus multiome sequencing of visceral adipose tissue from seven participants in the KOBS cohort

Visceral adipose tissue (VAT) biopsies were obtained from seven participants of the Kuopio OBesity Surgery Study (KOBS) cohort^23,24^ undergoing bariatric surgery. The participants of the KOBS cohort were recruited in the University of Eastern Finland and Kuopio University Hospital, Kuopio, Finland. All participants provided a written informed consent, and the KOBS study was approved by the Ethics Committee of the Northern Savo Hospital District, in accordance with the Declaration of Helsinki.

The seven VAT samples were processed using the 10x Single Cell Multiome ATAC + Gene Expression kit on a 10x Chromium controller. Briefly, we pooled seven VAT biopsies in equal ratios and isolated nuclei for droplet encapsulation, GEM formation, and cell barcoding. The fresh-frozen samples were first combined and then manually minced over dry ice. The tissue was then lysed in 500 μL of buffer (10 mM Tris-HCl, 10 mM NaCl, 3 mM MgCl_2_, 0.1% Tween-20, 0.1% IGEPAL CA-630, 0.01% Digitonin, 1% BSA, 1 mM DTT, and 1 U/μL RNase inhibitor) for 15 minutes on ice. The lysate was then mixed with 500 μl of wash buffer (10 mM Tris-HCl, 10 mM NaCl, 3 mM MgCl_2_, 1% BSA, 0.1% Tween-20, 1 mM DTT, and 1 U/μL RNase inhibitor) and filtered through a 70 μm FlowMi cell strainer. The nuclei were then centrifuged for 5 minutes at 500 x g at 4°C. This was followed by a resuspension in 1 ml wash buffer, filtering with a 40 μm FlowMi cell strainer, and another centrifugation at 500 x g for 5 minutes at 4°C. The resulting pellet was suspended in 30 μl of chilled nuclei buffer (1X Nuclei Buffer (10x Genomics), 1 mM DTT, and 1 U/μl RNase inhibitor). Concentration and quality were assessed using a Countess II Automated Cell Counter with trypan blue and DAPI staining. Joint single nucleus RNA and ATAC libraries were constructed with Single Cell Multiome ATAC + Gene Expression Reagent Kit (10x Genomics). Concentration and quality of cDNA and libraries were assessed using an Agilent Bioanalyzer. Finally, libraries were sequenced on an Illumina NovaSeq X Plus for the RNA and an Illumina NextSeq 500 for the ATAC, targeting 400 million reads in each modality. Reads were aligned with CellRanger Arc v2.0.2^25^ using GRCh38 and GENCODE v41^37^. Peak calls and read counts from CellRanger were then used for downstream analyses.

### Genotyping and imputation of KOBS participants

We genotyped the participants of the KOBS cohort using the Illumina Infinium Global Screening Array-24 v1. We used plink v1.9^39^ for basic filtering. Individuals with missingness > 2% were excluded, and we verified reported sex. We then removed unstranded or strand-ambiguous SNPs, monomorphic SNPs, SNPs with missingness > 2%, and SNPs with a Hardy-Weinberg Equilibrium (HWE) p-value < 1x10^-6^. The genotypes were then phased and imputed using the HRC reference panel r1.1 2016^40^ on the Michigan imputation server. SNPs with an allele mismatch to the reference were removed, phased with EAGLE v2.4^41^, and imputed with minimac4^42^. For demultiplexing, we removed monomorphic and kept biallelic genotyped and imputed SNPs with R^2^ > 0.99, resulting in 3,995,059 SNPs.

### Demultiplexing of simulated and VAT pooled multiome data

Demultiplexing of the simulated and VAT pooled multiome samples were performed in the same manner as follows. For ambimux, we used all inter + intra peak reads for the simulations as ambient fractions were kept constant for all genomic regions. We used intra peak reads for demultiplexing the VAT data with ambimux unless specified otherwise. For Vireo v0.5.8^15^, we first ran a pileup with cellSNP (Cellsnp-lite v1.2.3)^43^ using the BAM and VCF files. We set the UMI specifier to “UB” for the RNA runs and “None” for the ATAC, and required a minimum read count of 1 for variant pileup. We ran Vireo on the cellSNP pileup output using default parameters. For SouporCell v2.4^16^, we used ‘--skip_remap True’ and 200 restarts for both RNA and ATAC, ‘--no_umi True’ for the ATAC, and default settings otherwise. For Demuxlet v2, we ran the popscle implementation (https://github.com/statgen/popscle), where the pileup was generated first for input to Demuxlet. We set the minimum base quality score to 19, set an empty UMI tag with the ‘--tag-UMI’ parameter for the ATAC only, and used default parameters otherwise. Ambimux was run on all droplets without prior filtering, while Vireo, SouporCell, and Demuxlet were run on only the candidate set of droplets. These were defined as the known cell-containing barcodes for the simulations. For the VAT dataset, the candidates were defined as the 6,414 droplets with at least 100 RNA UMIs and 100 ATAC fragments from the 10x CellRanger barcode summary output. Comparisons between methods were restricted to these candidate droplets.

We called singlets and doublets and assigned donors based on the posterior probabilities. The posterior probabilities were calculated from the log probabilities for SouporCell and from the log likelihood difference between singlet and doublet for Demuxlet. We used a posterior probability of 0.90 to classify droplets as singlet or doublet and classified them as ambiguous otherwise. For ambimux, we also classified empty droplets using the same threshold. For donor assignment, cells identified as singlets were assigned to the donor with the highest likelihood score.

### Clustering and gene-peak link analysis of VAT pooled multiome data

To evaluate how ambient contamination affects the ability to detect gene-peak links, we clustered and separated the data by ambient fraction. First, we kept singlets identified by ambimux with a posterior probability greater than 0.90. To ensure enough coverage, we kept droplets with at least 200 RNA UMIs and 500 ATAC fragments. We processed and clustered the nuclei using Signac v1.10^38^, normalizing the RNA counts to sum to 1,000 and log transforming. For clustering and cell-type identification, we selected the top 2,000 variable genes using the variance-stabilizing transform, scaled the counts, and ran PCA. Then, we applied leiden clustering using a resolution of 0.5 on the top 50 PCs. Cell-types were assigned manually based on marker genes using ‘FindAllMarkers’. For visualization, we ran Uniform Manifold Approximation and Projection (UMAP) on the multimodal neighbor graph after processing the ATAC data with TFIDF and SVD on peaks with at least 5 counts.

We then separated droplets into low and high ambient groups ensuring equal coverage and cell-type representation. To do so, we first separated droplets by low ambient contamination (ambient fraction estimate less than 0.05 in both modalities) and high ambient contamination (ambient fraction estimate greater than 0.05 in both modalities). Then we binned droplets based on cell-type and coverage, using 10 equally-spaced bins based on log UMIs and log fragments. As the high ambient group had fewer nuclei, we sampled equal numbers of droplets without replacement from each group based on the bin distribution of the high ambient group. Finally, we ran the LinkPeaks Signac function^9,38^ on peaks within 500,000 base pairs of the target gene TSS and genes with at least 1 count in at least 10% of nuclei. P-values were corrected for multiple testing using FDR.

### Precision and recall curves

For the simulated data, we generated precision and recall curves against singlet posterior probability thresholds. Precision was defined as the fraction of singlets with a correct donor assignment over the number of called singlets. Recall was defined as the fraction of singlets with a correct donor assignment over the total number of singlets in the data.

### Estimation of ambient fractions with CellBender

We compared our ambient fraction estimation results with those from CellBender v0.2.2^18^, a method to remove ambient reads from the count matrix alone. We used the three simulated datasets of low, medium, and high contamination to have a ground truth metric for comparison. Evaluations were run for the RNA and ATAC modalities, although we note that CellBender was not explicitly designed for ATAC data. We set ‘--expected-cells 10000’ and ‘--total-droplets-included 11000’ to accurately reflect the number of droplets simulated and used default parameters otherwise. Ambient fraction estimates were then extracted from the ‘background_fraction’ field of the h5 output for the singlets.

## Data Availability

The visceral adipose tissue data (n=7) will be available under GEO accession number XXX upon acceptance. Due to privacy concerns, the genotype data (n=7) are available from the corresponding authors upon reasonable request and the data sharing involves a standard data sharing agreement.

## Code Availability

The ambimux software is available for download and use at https://github.com/marcalva/ambimux. The ambisim software used to generate the simulated data can be found at https://github.com/marcalva/ambisim.

## Acknowledgements

We would like to acknowledge and thank the participants of the KOBS cohort who participated in this study. We would like to acknowledge the Single Cell Genomics Core and Biocenter Finland for infrastructure support.

## Contributions

MA, EH, NZ, and PP conceived the project. MA designed the approach with contributions from TL, ER, ZC, OA, EH, CL, NZ, and PP. MA wrote the software. UTA, IS, TO, DK, VM, JP, MUK, and PP contributed towards generation of the visceral adipose tissue data. TL, STL, and AK performed data analyses and interpretation of the visceral adipose tissue data. DK, VM, JP, MUK, and PP collected cohort materials and data. MA, NZ, and PP wrote and contributed to the final manuscript. All authors read and approved of the final manuscript.

## Ethics approval and consent to participate

The KOBS study was approved by the Ethics Committee of the Northern Savo Hospital District. The study adhered to the principles outlined in the Declaration of Helsinki. All participants provided written informed consent.

