## Supplementary Note 1 and Figs. 1-9 for "Integrated ambient modeling and genetic demultiplexing of single-cell RNA+ATAC multiome experiments with Ambimux"

### Supplementary Note 1 Ambimux

#### 1.1 Model description

We consider a single-cell dataset of $N$ pooled genetically distinct donors. There are $D$ droplets, and for each droplet $d$, there are $M_d$ molecules (reads). Furthermore, $V_{dm}$ specifies the number of variants overlapped by molecule $m$. Droplets may be contaminated with cell-free RNA or DNA that do not originate from the cell. Variants are detected from RNA and ATAC reads at genotyped/imputed sites.

We use integer donor indices $i, j \in \{1, \ldots, N\}$ and distinguish donor indices from donor sets. Let $\mathcal{S}_0 = \{\varnothing\}$, $\mathcal{S}_1 = \{\{i\} : i = 1, \ldots, N\}$, and $\mathcal{S}_2 = \{\{i, j\} : 1 \leq i < j \leq N\}$. In particular, the third index $s$ in $\alpha_{dhs}$ denotes a realized donor set $s \in \mathcal{S}_h$.

We define the following variables:

- $H_d \in \{0, 1, 2\}$: number of nuclei in droplet $d$.
- $S_d \subseteq \{1, \ldots, N\}$: set of donor indices present in droplet $d$ (excluding ambient), with $S_d \in \mathcal{S}_{H_d}$.
- $T_{dm} \in \mathcal{T} = \{a, 1, \ldots, N\}$: source of molecule $m$ in droplet $d$; $a$ denotes the ambient pool and $1, \ldots, N$ denote donor indices.
- $B_{dmv} \in \{0, 1\}$: observed base call at variant $v$ in molecule $m$ in droplet $d$.
- $E_{dmv} \in \{0, 1\}$: sequencing error indicator for variant $v$ in molecule $m$ in droplet $d$.

The probability of observing a reference or alternate allele base call $b$ given source $t \in \mathcal{T}$ and sequencing error $E \in \{0, 1\}$ is a Bernoulli

$$p(B_{dmv} = b,\, E_{dmv} = 0 \mid T_{dm} = t) = (1 - \tau_{dmv})\, \gamma_{tv}^{b} (1 - \gamma_{tv})^{1-b},$$
$$p(B_{dmv} = b,\, E_{dmv} = 1 \mid T_{dm} = t) = \frac{\tau_{dmv}}{2},$$

which marginalizes to

$$p(B_{dmv} = b \mid T_{dm} = t) = (1 - \tau_{dmv})\, \gamma_{tv}^{b} (1 - \gamma_{tv})^{1-b} + \frac{\tau_{dmv}}{2}.$$

Here $\tau_{dmv}$ is the sequencing error probability derived from Phred quality scores. The genotype for variant $v$ of source $t$ is given by $\gamma_{tv}$. The ambient genotype is an average of the donor allele frequencies

$$\gamma_{av} = \sum_{i=1}^{N} \pi_{ai}\, \gamma_{iv}, \qquad \sum_{i=1}^{N} \pi_{ai} = 1,\ \pi_{ai} \geq 0, \tag{S1}$$

weighted by their composition in the ambient pool $\pi_a$.

We treat multiple variants in a molecule independently for simplicity, and the probability of observing $V_{dm}$ base calls in a molecule becomes

$$p(B_{dm} \mid T_{dm} = t) = \prod_{v=1}^{V_{dm}} p(B_{dmv} \mid T_{dm} = t). \tag{S2}$$

A molecule $m$ in droplet $d$ can originate from the ambient pool or a donor. Using state-specific contamination parameters $\alpha_{dhs} \in [0, 1]$ for droplet $d$ in state $(H_d = h, S_d = s)$, the categorical probabilities are

$$p(T_{dm} = t \mid H_d = h, S_d = s; \alpha_{dhs}) = \begin{cases} 1, & h = 0,\ t = a, \\ \alpha_{d,1,\{i\}}, & h = 1,\ s = \{i\},\ t = a, \\ 1 - \alpha_{d,1,\{i\}}, & h = 1,\ s = \{i\},\ t = i, \\ \alpha_{d,2,\{i,j\}}, & h = 2,\ s = \{i, j\},\ t = a, \\ \frac{1 - \alpha_{d,2,\{i,j\}}}{2}, & h = 2,\ s = \{i, j\},\ t \in \{i, j\}, \\ 0, & \text{otherwise}, \end{cases} \tag{S3}$$

with $1 \le i < j \le N$ and $\alpha_{dhs}$ specifying the state-specific droplet contamination fraction.

We place a categorical prior on the droplet nuclei parameterized by $\lambda$:

$$p(H_d = i; \lambda) = \lambda_i, \quad i \in \{0, 1, 2\}, \qquad \sum_{i=0}^{2} \lambda_i = 1,\ \lambda_i \ge 0.$$

Conditional on $H_d$, the donor set $S_d$ is drawn as follows. For empty droplets ($H_d = 0$) the set is empty. For singlets ($H_d = 1$), a single donor is drawn from $\pi_c$, the donor composition in the nuclei (as opposed to ambient). For doublets, an unordered pair of distinct donors $\{i, j\}$ with $i < j$ is chosen with probability proportional to $\pi_{ci}\pi_{cj}$ and normalized over all pairs. Formally,

$$\begin{aligned} p(S_d = \varnothing \mid H_d = 0; \pi_c) &= 1, \\ p(S_d = \{i\} \mid H_d = 1; \pi_c) &= \pi_{ci}, \quad i = 1, \dots, N, \\ p(S_d = \{i, j\} \mid H_d = 2; \pi_c) &= \frac{\pi_{ci}\, \pi_{cj}}{\sum_{1 \le p < q \le N} \pi_{cp}\, \pi_{cq}}, \quad 1 \le i < j \le N. \end{aligned}$$

The likelihood for droplet $d$ factorizes as

$$p(X_d \mid Z_d; \Theta) = p(H_d; \lambda)\, p(S_d \mid H_d; \pi_c) \prod_{m=1}^{M_d} p(T_{dm} \mid H_d, S_d; \alpha_{d,H_d,S_d}) \prod_{v=1}^{V_{dm}} p(B_{dmv} \mid T_{dm}; \gamma, \tau, \pi_a),$$

where

- $X_d = \{B_{dmv} : m = 1, \dots, M_d,\ v = 1, \dots, V_{dm}\}$ (observed data),
- $Z_d = \{H_d, S_d, T_{d1}, \dots, T_{dM_d}\}$ (latent variables),
- $\Theta = \{\lambda, \pi_c, \pi_a, \{\alpha_{dhs}\}, \gamma, \tau\}$ (parameters).

For EM, the expected log-likelihood with respect to the posterior of the latent variables is

$$Q(\Theta \mid \Theta^{(t)}) = \mathbb{E}_{P(Z|X,\Theta^{(t)})}\big[\log L(\Theta \mid X, Z)\big].$$

#### 1.2 Optimization of $\pi_a$

We define the donor composition in the ambient pool as $\pi_a = (\pi_{a1}, \dots, \pi_{aN})$ with $\sum_{i=1}^{N} \pi_{ai} = 1$ and $\pi_{ai} \ge 0$. This allows us to more accurately model allele frequencies of the ambient pool for better estimation of droplet contaminations. To estimate $\pi_a$, we use gradient ascent on the log likelihood, which allows for faster convergence than optimization of the expected log likelihood.

The likelihood depends on the parameter $\pi_a$ through $\gamma_{av}$ in the probability term modeling a base call in an ambient molecule. Specifically, for a molecule $(d, m)$ and variant $v$ with ambient source $T_{dm} = a$, the probability of the observed base call $B_{dmv}$ is

$$p(B_{dmv} \,|\, T_{dm} = a) = (1 - \tau_{dmv})\, A(\gamma_{av}) + \frac{\tau_{dmv}}{2}, \quad A(u) := u^{B_{dmv}} (1 - u)^{1 - B_{dmv}},$$

where $\gamma_{av} = \sum \pi_{ai}\gamma_{iv}$ is given by equation (S1). The log-likelihood terms that depend on $\pi_a$ (through $\gamma_{av}$) are those with $T_{dm} = a$:

$$\ell(\pi_a) = \sum_{d,m,v:\, T_{dm}=a} \log\left[(1 - \tau_{dmv})\, A(\gamma_{av}) + \frac{\tau_{dmv}}{2}\right].$$

Abbreviating $\tau := \tau_{dmv}$ and $B := B_{dmv}$, the derivative with respect to $\pi_{ai}$ is given by

$$\frac{\partial}{\partial \pi_{ai}} \log p(B_{dmv} \,|\, T_{dm} = a) = \frac{(1 - \tau)\, A'(\gamma_{av})}{(1 - \tau)\, A(\gamma_{av}) + \frac{\tau}{2}} \, \frac{\partial \gamma_{av}}{\partial \pi_{ai}}.$$

Because $\gamma_{av}$ depends linearly on $\pi_{ai}$, we have

$$\frac{\partial \gamma_{av}}{\partial \pi_{ai}} = \gamma_{iv}.$$

Using $A'(u) = A(u)\,\dfrac{B-u}{u(1-u)}$ evaluated at $u = \gamma_{av}$, the per-term gradient is

$$\frac{\partial}{\partial \pi_{ai}} \log p(B_{dmv} \,|\, T_{dm} = a) = (1-\tau)\,\gamma_{iv}\,\frac{A(\gamma_{av})}{(1-\tau)A(\gamma_{av}) + \frac{\tau}{2}}\,\frac{B-\gamma_{av}}{\gamma_{av}(1-\gamma_{av})}.$$

Summing over all ambient-source terms yields the total gradient

$$\frac{\partial\,\ell(\pi_a)}{\partial \pi_{ai}} = \sum_{d,m,v:\,T_{dm}=a} \frac{(1-\tau_{dmv})\,\gamma_{iv}}{\gamma_{av}(1-\gamma_{av})}\,\frac{\gamma_{av}^{B_{dmv}}(1-\gamma_{av})^{1-B_{dmv}}\,(B_{dmv}-\gamma_{av})}{(1-\tau_{dmv})\,\gamma_{av}^{B_{dmv}}(1-\gamma_{av})^{1-B_{dmv}} + \frac{\tau_{dmv}}{2}}.$$

We update $\pi_a$ via projected gradient ascent:

$$\pi_a^{(t+1)} = \Pi_\Delta\big(\,\pi_a^{(t)} + \eta\,\nabla_{\pi_a}\,\ell(\pi_a^{(t)})\,\big),$$

where $\Pi_\Delta$ denotes a Euclidean projection onto the probability simplex (Chen and Ye 2011). We use a fixed step size $\eta = 1/|V_a|$, with $|V_a|$ the number of variants across molecules in ambient droplets. The updates iterate until $||\pi_a^{(t+1)} - \pi_a^{(t)}||_2/||\pi_a^{(t)}||_2 < \varepsilon$ with $\varepsilon = 10^{-6}$. For computational efficiency, we only use fixed empty droplets and set $\pi_a$ once before EM.

#### 1.3 Optimization of $\lambda$

The parameters $\lambda = \{\lambda_0, \lambda_1, \lambda_2\}$ represent the prior probabilities for the number of nuclei in each droplet, where $\lambda_i = p(H_d = i)$ for $i \in \{0, 1, 2\}$. We maximize $\lambda$ using the EM algorithm by extracting terms from the expected log-likelihood $Q$ that depend on these parameters:

$$Q(\Theta \mid \Theta^{(t)}) \supset \sum_{d\in\mathcal{D}} \mathbb{E}_{P(Z|X,\Theta^{(t)})}\big[\log p(H_d \mid \lambda)\big] = \sum_{d\in\mathcal{D}}\sum_{i=0}^{2} p(H_d = i \mid X; \Theta^{(t)}) \log \lambda_i.$$

To maximize this subject to the constraint $\sum_{i=0}^{2} \lambda_i = 1$, we use Lagrange multipliers. The Lagrangian is:

$$\mathcal{L} = \sum_{d\in\mathcal{D}}\sum_{i=0}^{2} p(H_d = i \mid X; \Theta^{(t)}) \log \lambda_i - \mu\left(\sum_{i=0}^{2} \lambda_i - 1\right).$$

Taking the derivative with respect to $\lambda_i$ and setting to zero:

$$\frac{\partial \mathcal{L}}{\partial \lambda_i} = \frac{\sum_{d\in\mathcal{D}} p(H_d = i \mid X; \Theta^{(t)})}{\lambda_i} - \mu = 0.$$

This gives

$$\lambda_i = \frac{\sum_{d\in\mathcal{D}} p(H_d = i \mid X; \Theta^{(t)})}{\mu}.$$

Using the constraint to solve for $\mu$:

$$\mu = \sum_{i=0}^{2}\sum_{d\in\mathcal{D}} p(H_d = i \mid X; \Theta^{(t)}) = |\mathcal{D}|.$$

Therefore, the M-step update for $\lambda$ is:

$$\hat{\lambda}_i = \frac{\sum_{d\in\mathcal{D}} p(H_d = i \mid X; \Theta^{(t)})}{|\mathcal{D}|}.$$

This represents the empirical frequency of droplets assigned to each nuclei count, weighted by the posterior probabilities from the E-step.

#### 1.4 Optimization of $\pi_c$

The parameters $\pi_c = (\pi_{c1}, \dots, \pi_{cN})$ are the prior probabilities that a nucleus belongs to each donor, with $\sum_{i=1}^{N} \pi_{ci} = 1$ and $\pi_{ci} \geq 0$.

The part of the expected log-likelihood $Q(\Theta \mid \Theta^{(t)})$ that depends on $\pi_c$ collects contributions from singlets and doublets:

$$\begin{aligned} Q(\Theta \mid \Theta^{(t)}) \supset & \sum_{d \in \mathcal{D}} \sum_{i=1}^{N} p\big(H_d = 1,\, S_d = \{i\} \mid X; \Theta^{(t)}\big) \log \pi_{ci} \\ & + \sum_{d \in \mathcal{D}} \sum_{1 \leq i < j \leq N} p\big(H_d = 2,\, S_d = \{i, j\} \mid X; \Theta^{(t)}\big) \, (\log \pi_{ci} + \log \pi_{cj}). \end{aligned}$$

Define the posterior-weighted expected nucleus count for donor $i$ as

$$C_i = \sum_{d \in \mathcal{D}} p\big(H_d = 1,\, S_d = \{i\} \mid X; \Theta^{(t)}\big) + \sum_{d \in \mathcal{D}} \sum_{\substack{j=1 \\ j \neq i}}^{N} p\big(H_d = 2,\, S_d = \{i, j\} \mid X; \Theta^{(t)}\big),$$

so that the $\pi_c$-dependent objective simplifies to

$$Q_{\pi_c}(\pi_c) = \sum_{i=1}^{N} C_i \, \log \pi_{ci}.$$

To maximize $Q_{\pi_c}$ under the simplex constraint, we introduce a Lagrange multiplier $\mu$ and form the Lagrangian

$$\mathcal{L}(\pi_c, \mu) = \sum_{i=1}^{N} C_i \, \log \pi_{ci} - \mu \left( \sum_{i=1}^{N} \pi_{ci} - 1 \right).$$

Setting the derivatives to zero yields

$$\frac{\partial \mathcal{L}}{\partial \pi_{ci}} = \frac{C_i}{\pi_{ci}} - \mu = 0 \quad \Rightarrow \quad \pi_{ci} = \frac{C_i}{\mu}, \qquad i = 1, \dots, N.$$

Enforcing $\sum_i \pi_{ci} = 1$ gives $\mu = \sum_{i=1}^{N} C_i$ and the M-step update

$$\hat{\pi}_{ci} = \frac{C_i}{\sum_{j=1}^{N} C_j}.$$

This sets $\pi_{ci}$ to the posterior-weighted empirical frequency of donor $i$ across nuclei: each singlet contributes one nucleus for its donor, and each doublet contributes one nucleus for each of its two donors.

#### 1.5 Optimization of $\alpha_{dhs}$

The modality-specific contamination parameters $\alpha_{dhs}$ represent the proportion of ambient material in droplet $d$ when it contains $h$ nuclei with donor composition $s$. Each valid combination of $(H_d, S_d)$ has its own contamination parameter, constrained to $[0, 1]$. We optimize these parameters using Newton–Raphson on the log marginal likelihood, treating each $\alpha_{dhs}$ independently. This allows for much faster convergence than optimization of the expected log likelihood during EM. We first derive the likelihood and then the first and second derivatives with respect to $\alpha_{dhs}$.

For a droplet $d$ with state $(H_d = h, S_d = s)$, where $s$ denotes the donor set $s \in \mathcal{S}_{\mathcal{H}_\Gamma}$, the marginal likelihood is

$$\begin{aligned} L(\alpha_{dhs}) =& p(X_d \mid H_d = h, S_d = s; \alpha_{dhs}) \\ =& \prod_{m=1}^{M_d} \sum_{t \in \mathcal{T}_{hs}} p(T_{dm} = t \mid H_d = h, S_d = s; \alpha_{dhs}) \, p(B_{dm} \mid T_{dm} = t), \end{aligned}$$

where $\mathcal{T}_{hs} = \{a\} \cup s$ is the set of possible molecule sources (ambient plus the donors in the set $s$). The probability of $T_{dm}$ is given by equation (S3) and the probability of $B_{dmv}$ is given by equation (S2).

For empty droplets ($H_d = 0$), we fix $\alpha_{d,0,\varnothing} = 1$ and no optimization is needed.

Next, we expand the sum for singlets and doublets separately and include the term $\alpha_{dhs}$. For a singlet of donor $i$ ($H_d = 1, S_d = \{i\}$), the marginal likelihood is

$$\begin{aligned} L(\alpha_{d,1,\{i\}}) &= \prod_{m=1}^{M_d} \Big[\alpha_{d,1,\{i\}}\, p(B_{dm} \mid T_{dm} = a) \\ &\quad + (1 - \alpha_{d,1,\{i\}})\, p(B_{dm} \mid T_{dm} = i)\Big] \\ &= \prod_{m=1}^{M_d} \Phi_m(\alpha_{d,1,\{i\}}). \end{aligned}$$

For a doublet of donors $i$ and $j$, ($H_d = 2, S_d = \{i, j\}$), the marginal likelihood is

$$\begin{aligned} L(\alpha_{d,2,\{i,j\}}) &= \prod_{m=1}^{M_d} \Big[\alpha_{d,2,\{i,j\}}\, p(B_{dm} \mid T_{dm} = a) \\ &\quad + \frac{1 - \alpha_{d,2,\{i,j\}}}{2} \big(p(B_{dm} \mid T_{dm} = i) + p(B_{dm} \mid T_{dm} = j)\big)\Big] \\ &= \prod_{m=1}^{M_d} \Psi_m(\alpha_{d,2,\{i,j\}}). \end{aligned}$$

We maximize the log of the marginal likelihood after adding a log beta prior to prevent boundary solutions and avoid collapse, given by

$$R(\alpha; \omega, \beta) = (\omega\beta - 1) \log \alpha + (\omega(1 - \beta) - 1) \log(1 - \alpha),$$

where $\beta$ represents the expected ambient contamination level and $\omega$ controls regularization strength. Effectively, this adds $\omega\beta$ ambient reads and $\omega(1 - \beta)$ donor reads.

Using the shorthand $\alpha = \alpha_{dhs}$ for readability, the regularized log marginal likelihoods are

$$\ell(\alpha) = \sum_{m=1}^{M_d} \log \Phi_m(\alpha) + R(\alpha; \omega, \beta),$$

$$\ell(\alpha) = \sum_{m=1}^{M_d} \log \Psi_m(\alpha) + R(\alpha; \omega, \beta),$$

for singlets and doublets, respectively.

The first and second derivatives for singlets are

$$\ell'(\alpha) = \sum_{m=1}^{M_d} \frac{\Phi'_m(\alpha)}{\Phi_m(\alpha)} + R'(\alpha; \omega, \beta),$$

$$\ell''(\alpha) = -\sum_{m=1}^{M_d} \frac{\Phi'_m(\alpha)^2}{\Phi_m(\alpha)^2} + R''(\alpha; \omega, \beta),$$

while the derivatives for doublets are

$$\ell'(\alpha) = \sum_{m=1}^{M_d} \frac{\Psi'_m(\alpha)}{\Psi_m(\alpha)} + R'(\alpha; \omega, \beta),$$
$$\ell''(\alpha) = -\sum_{m=1}^{M_d} \frac{\Psi'_m(\alpha)^2}{\Psi_m(\alpha)^2} + R''(\alpha; \omega, \beta),$$

where

$$\Phi'_m(\alpha) = p(B_{dm} \mid T_{dm} = a) - p(B_{dm} \mid T_{dm} = i),$$
$$\Psi'_m(\alpha) = p(B_{dm} \mid T_{dm} = a) - \frac{1}{2}p(B_{dm} \mid T_{dm} = i) - \frac{1}{2}p(B_{dm} \mid T_{dm} = j),$$
$$R'(\alpha; \omega, \beta) = \frac{\omega\beta - 1}{\alpha} - \frac{\omega(1-\beta) - 1}{1-\alpha},$$
$$R''(\alpha; \omega, \beta) = -\frac{\omega\beta - 1}{\alpha^2} - \frac{\omega(1-\beta) - 1}{(1-\alpha)^2}.$$

We initialize all $\alpha_{dhs} = 0.5$, $\beta = 0.1$, and set $\omega = 10^{-8}$ for weak regularization. Each $\alpha_{dhs}$ is optimized with Newton–Raphson until the absolute change in paramater value is $< \varepsilon = 10^{-6}$, or for a maximum of 500 iterations. The weight $\omega$ is kept fixed while $\beta$ is learned from the data. The prior $\beta$ is updated to the average $\alpha_{dhs}$ in singlets weighted by the posterior probability $P(H_d = 1, S_d \mid X_d; \Theta^{(t)})$. This optimization occurs within each M-step of the EM algorithm.

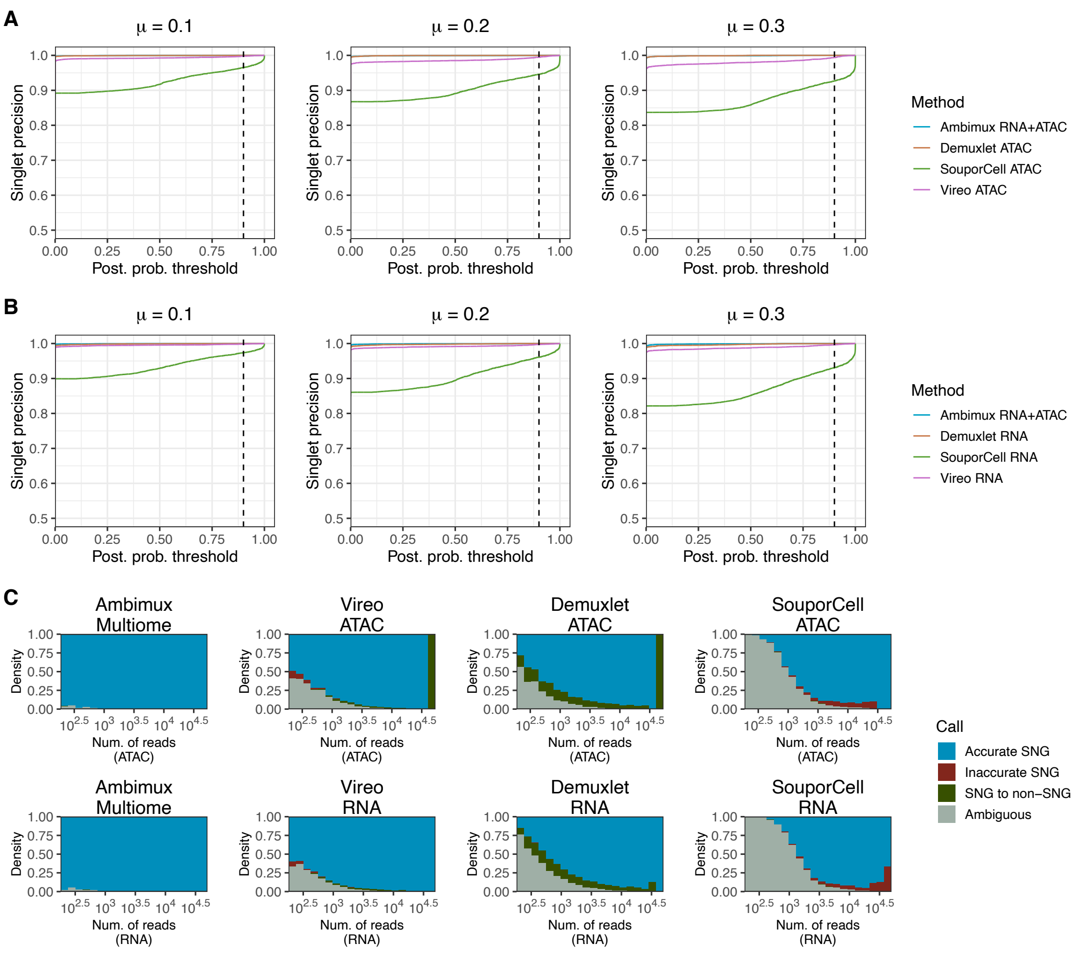


**Fig S1: Singlet assignment precision of demultiplexing simulated multiome datasets.** **A, B** Precision of singlet assignment for three simulated multiome datasets with low (mean 0.1), medium (mean 0.2), and high (mean 0.3) ambient contamination. Precision is defined as the proportion of called singlets with a correct donor assignment. The precision curves compare ambimux run on multimodal data with competing methods run on ATAC (**A**) and RNA (**B**). **C** Stacked bar plots showing the proportion of ground truth singlets that are correctly assigned, assigned to an incorrect donor, classified as a non-singlet, and unassigned (ambiguous call) binned by number of reads. The plots compare multimodal ambimux against competing methods run on RNA (top) and ATAC (bottom).

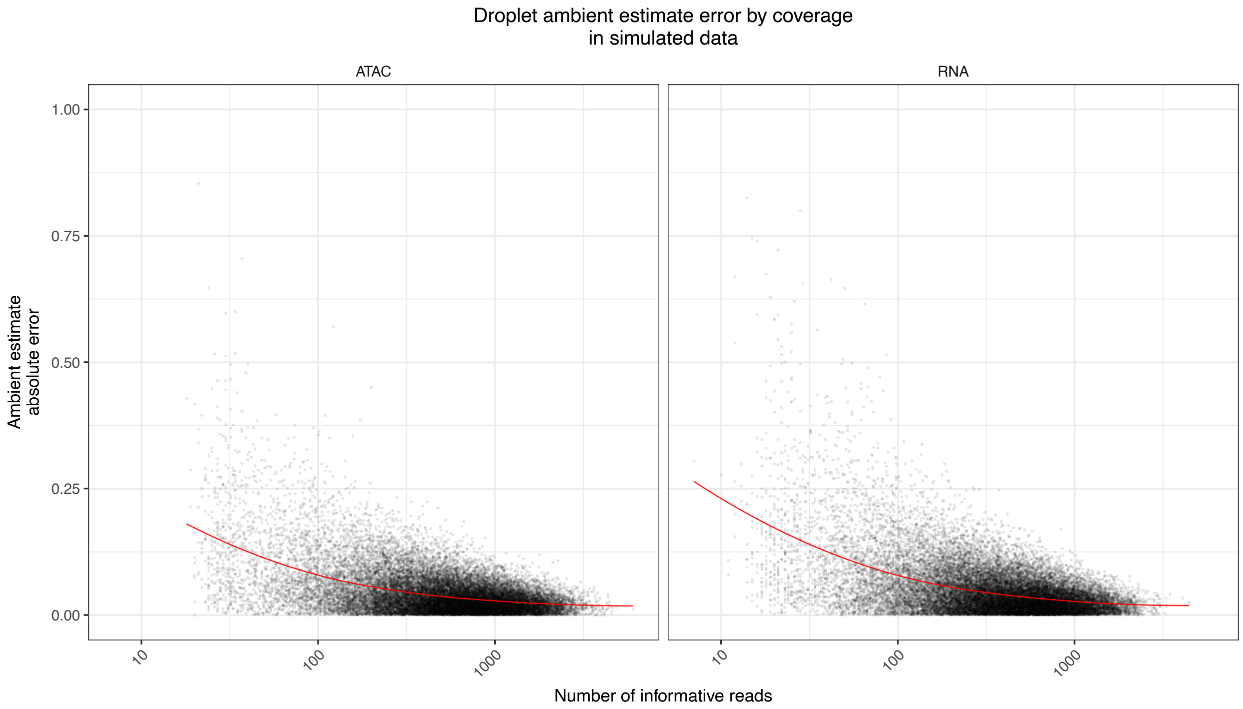


**Fig S2: Ambient estimation accuracy improves with increased coverage.** Relationship between the number of informative reads in a droplet against the absolute error in ambient fraction estimation. Informative reads are defined as those that overlap a variant used for demultiplexing.

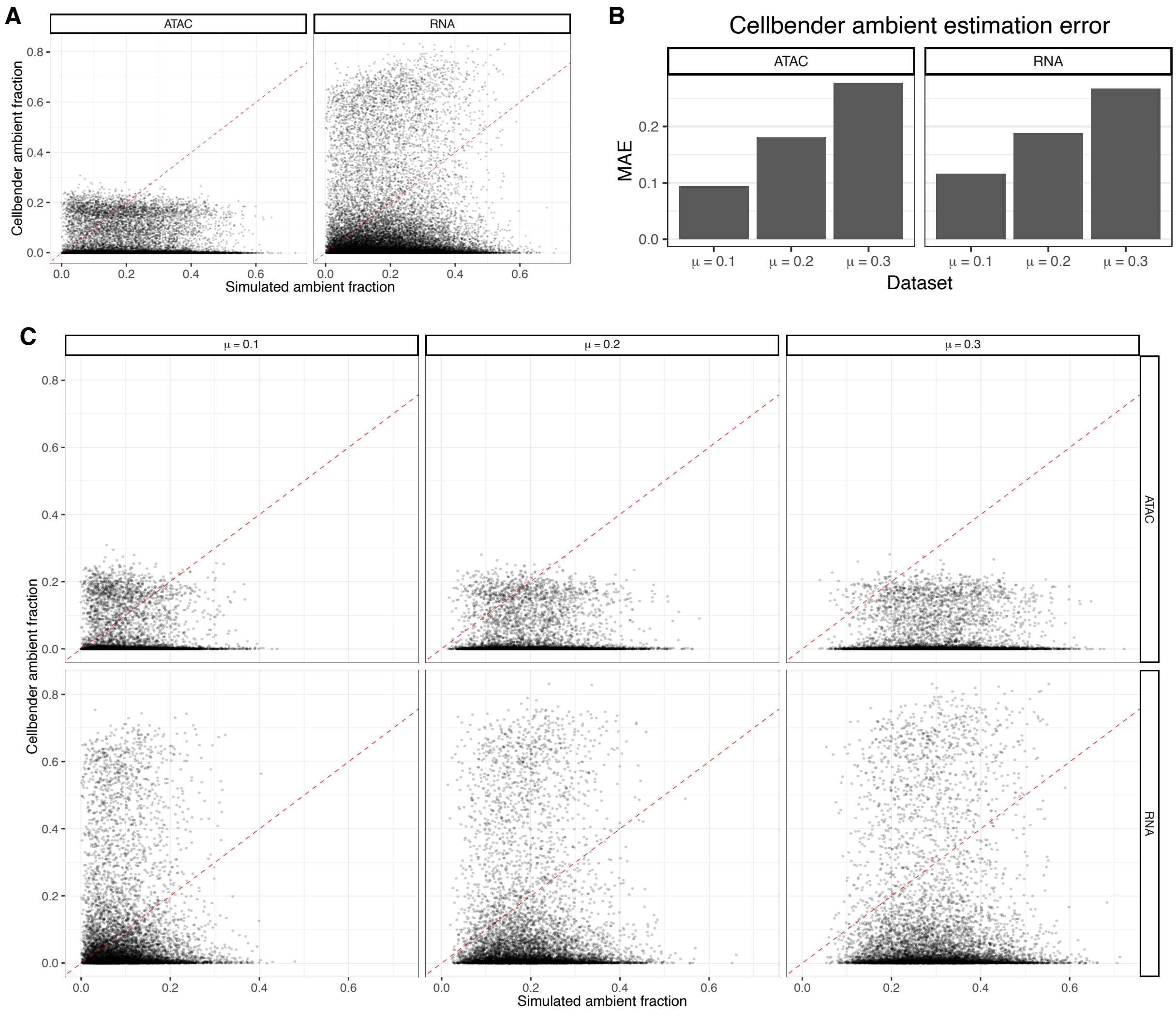


**Fig S3: Ambient fraction estimates from CellBender on simulated single-cell multiome data. A** Correlation between CellBender ambient estimates and simulated ground truth ambient estimates after combining results for the low, medium, and high ambient simulations in ATAC and RNA. **B** The mean absolute error (MAE) of ambient estimates in the low, medium, and high ambient datasets for ATAC and RNA. **C** Expanded correlation plot that shows the ambient estimates vs. simulated ground truth in each of the three datasets for ATAC and RNA.

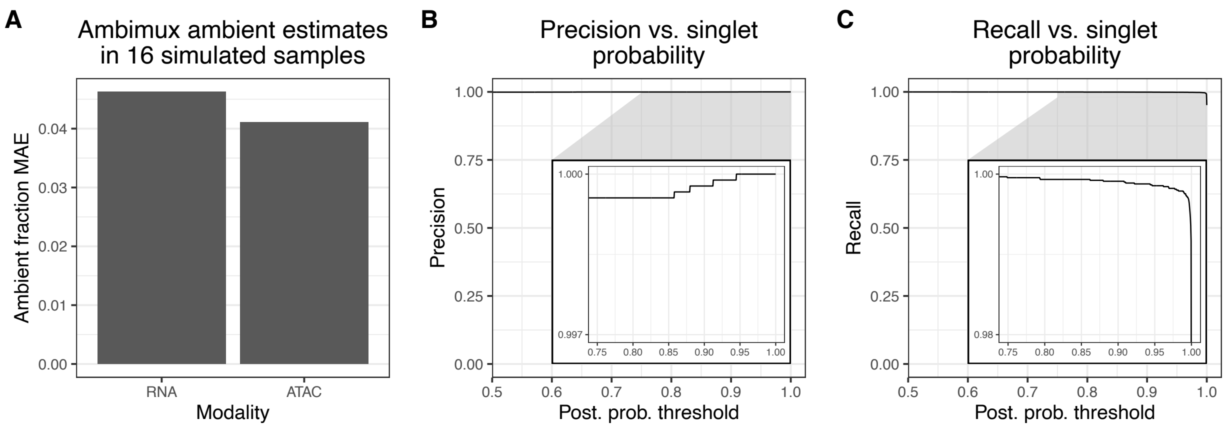


**Fig. S4: Ambimux maintains high accuracy in a multiome simulation of varying donor proportions. A** The mean absolute error (MAE) of ambient estimates in a simulated dataset of 16 donors with donor proportions independently varying in the background and nuclei pool. **B, C** Precision (**B**) and recall (**C**) curves against the posterior probability threshold for the same simulated dataset as in (**A**).

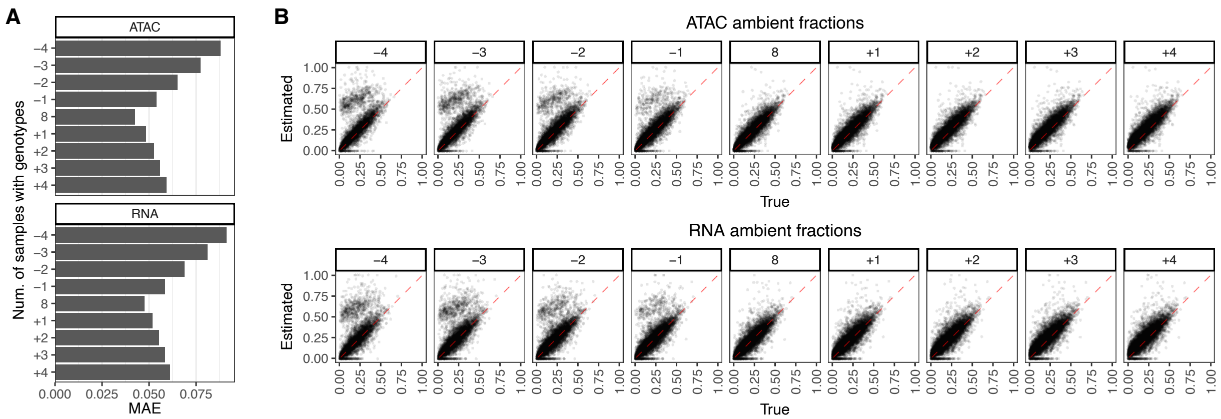


**Fig. S5: Effects of artificially adding or removing genotype donors on ambimux ambient fraction estimation.** The accuracy of ambient estimation slightly worsens when genotypes are added or removed. Simulated data were generated for eight donors, but genotypes were either removed (negative numbers) or added (positive numbers). **A** The mean absolute error (MAE) of the ATAC (top) and RNA (bottom) ambient estimates from ambimux when demultiplexing after removing genotype samples (negative numbers) and adding genotype samples (positive numbers). **B** Correlation of true vs. estimated ambient fractions for the ATAC (top) and RNA (bottom).

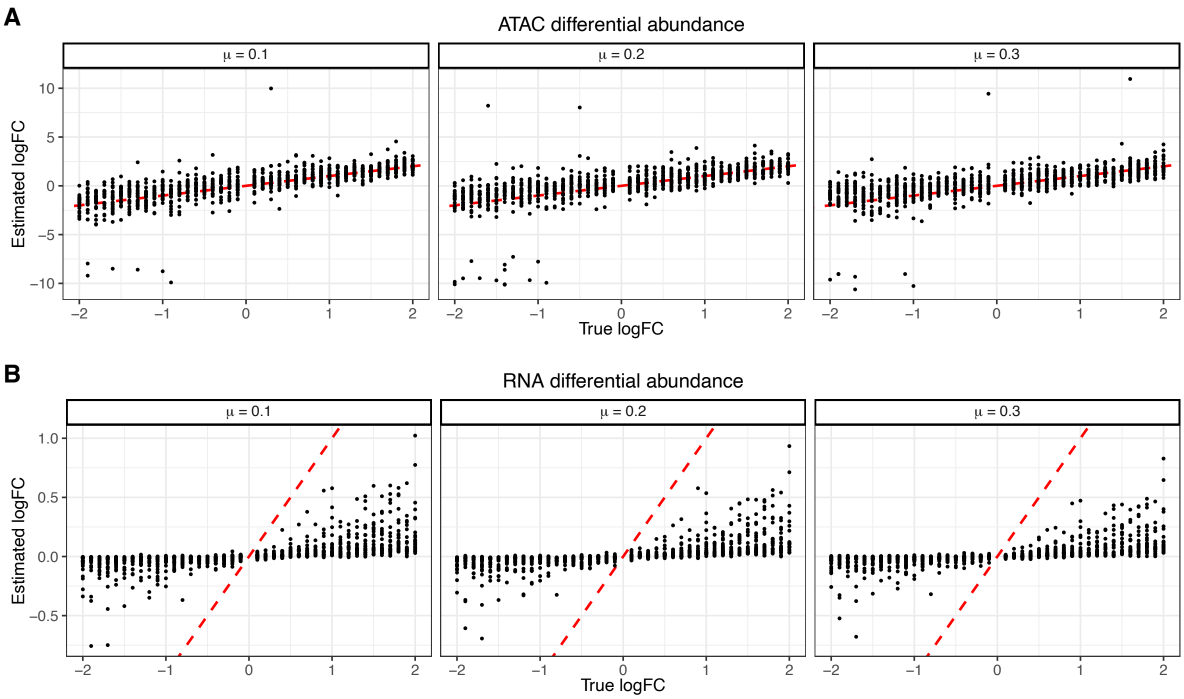


**Fig. S6: Differential abundance of genes and peaks with varying levels of ambient contamination. A, B** Correlation of estimated vs. true log fold-changes for ATAC (**A**) and RNA (**B**) in a combined simulated multiome dataset of low (mean 10%), medium (mean 20%), and high (mean 30%) ambient contamination fraction. A pool of eight donors were simulated, of which four contained a disease cell subtype with 1,000 genes and 1,000 peaks differentially abundant with varying log fold-changes.

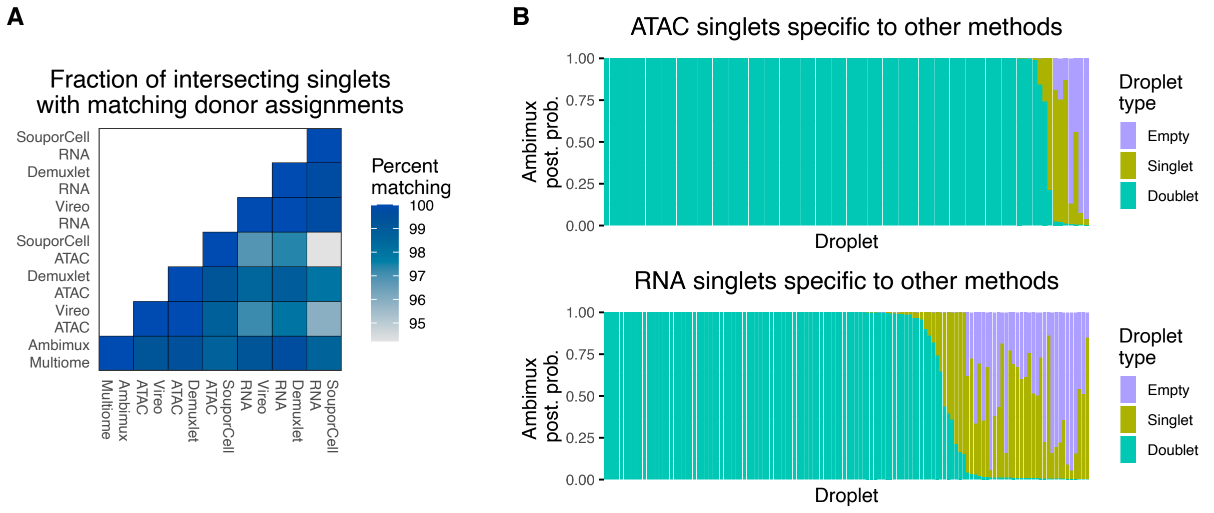


**Fig. S7: Comparison of demultiplexing approaches in visceral adipose tissue of samples pooled. A** Proportion of singlets with a concordant donor assignment between methods in visceral adipose tissue pool of seven donors. Among droplets classified as singlets in two methods, the plot shows the percent that are assigned to the same donor. **B** Ambimux droplet type probability for droplets classified as a singlet in SoupOrCell, Demuxlet, or Vireo but not in ambimux. Ambimux was run on the multiome data, and singlets were taken from methods run in the ATAC (top) or in the RNA (bottom) modalities. The bar plot shows that most singlets specific to other methods are classified as doublets by ambimux.

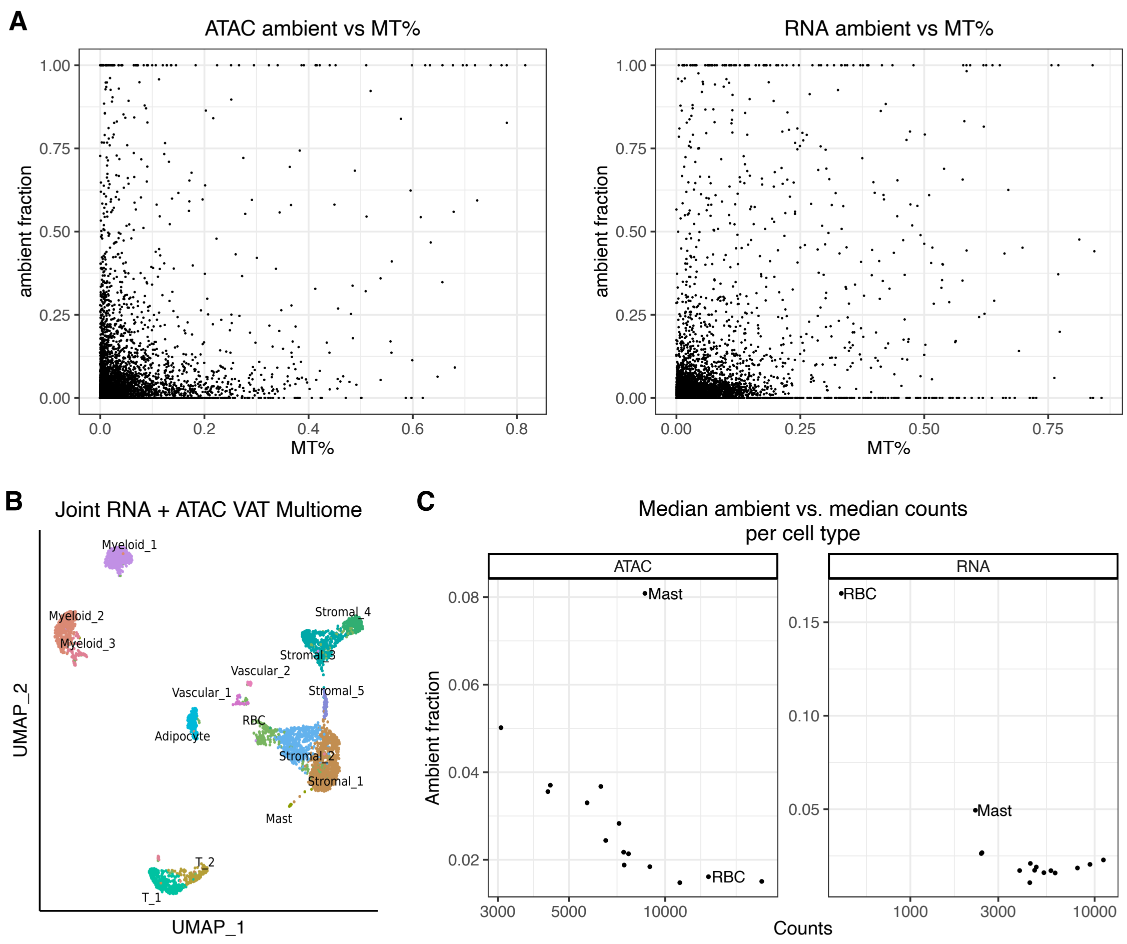


**Fig. S8: Correlation of read-based metrics with ambient fraction estimates in visceral adipose tissue multiome. A** Modality-specific ambient fraction estimates from ambimux show a low positive correlation with percent of mitochondrial reads in ATAC (Spearman rho = 0.20) and RNA (Spearman rho = 0.25). **B** UMAP visualization of ambimux singlets colored by cell-type using joint RNA+ATAC clustering. **C** Correlation between cell-type median ATAC fragment (left) or RNA UMI (right) count and mean ambient fraction estimate, showing low-coverage droplets tend to have higher background fractions. Mast and red blood cells (RBC) are labelled to show the lowest coverage cell-types in RNA.

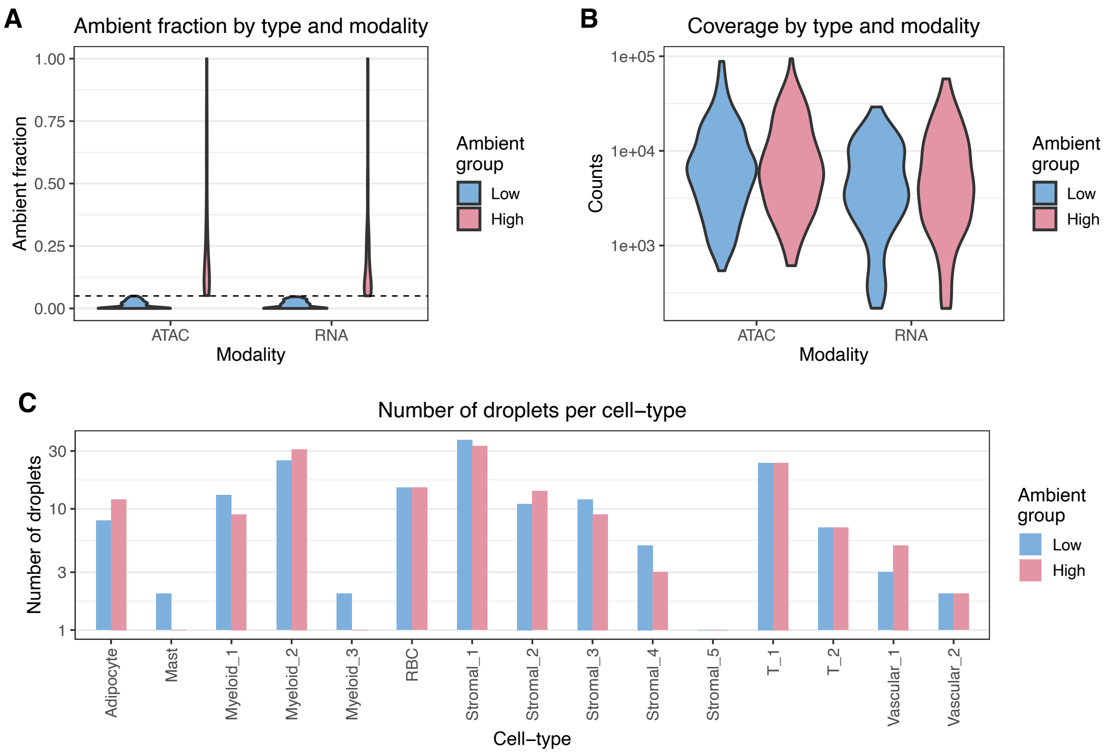


**Fig. S9: Controlled separation of low-ambient and high-ambient fractions droplets in visceral adipose tissue multiome.** Droplets from the visceral adipose tissue dataset were separated into low-ambient (< 5% background percent in both modalities) and high-ambient (> 5% background percent in both modalities) groups while controlling for read coverage and cell-type representation. **A** Ambient fraction plot showing the cutoff used to group droplets and their similar distribution in both modalities. **B** Distribution of fragment and UMI counts for ATAC and RNA modalities, respectively, grouped by low vs. high ambient fraction. **C** Number of droplets per cell-type grouped by low vs. high ambient group, showing similar representation between the two groups.
